# Testing for dependence on tree structures

**DOI:** 10.1101/622811

**Authors:** Merle Behr, M. Azim Ansari, Axel Munk, Chris Holmes

## Abstract

Tree structures, showing hierarchical relationships and the latent structures between samples, are ubiquitous in genomic and biomedical sciences. A common question in many studies is whether there is an association between a response variable measured on each sample and the latent group structure represented by some given tree. Currently this is addressed on an ad hoc basis, usually requiring the user to decide on an appropriate number of clusters to prune out of the tree to be tested against the response variable. Here we present a statistical method with statistical guarantees that tests for association between the response variable and a fixed tree structure across all levels of the tree hierarchy with high power, while accounting for the overall false positive error rate. This enhances the robustness and reproducibility of such findings.

**Significance Statement:** Tree like structures are abundant in the empirical sciences as they can summarize high dimensional data and show latent structure among many samples in a single framework. Prominent examples include phylogenetic trees or hierarchical clustering derived from genetic data. Currently users employ ad hoc methods to test for association between a given tree and a response variable, which reduces reproducibility and robustness. In this paper, we introduce treeSeg, a simple to use and widely applicable methodology with high power for testing between all levels of hierarchy for a given tree and the response while accounting for the overall false positive rate. Our method allows for precise uncertainty quantification and therefore increases interpretability and reproducibility of such studies across many fields of science.

---

In the era of big data where quantifying the relationship between samples is difficult, tree structures are commonly used to summarize and visualize the relationship between samples and to capture latent structure. The hierarchical nature of trees allows the relationships between all samples to be viewed in a single framework and this has lead to their widespread usage in genomics and biomedical science. Examples are phylogenetic trees built from genetic data, hierarchical clustering based on distance measures of features of interest (for example gene expression data with thousands of markers measured in each sample), evolution of human languages, and more broadly in machine learning where clustering and unsupervised learning is a fundamental task (1–7).

Often samples have additional response measurements *y*_*i*_ (e.g., phenotypes) and a common question is whether there is a relation between the sample’s latent group structure captured by the tree *T* and the outcome of interest *y*_*i*_, i.e., whether the distribution of *y*_*i*_ depends on its relative location among the leaves of the tree *T*. Testing for all possible combinations of groupings on the tree is practically impossible as it grows exponentially with sample size. Currently users typically decide on the number of clusters on an ad hoc basis (e.g., after plotting the response measurement on the leaves of the tree and deciding visually which clusters to choose). These clusters are then tested for association with the outcome of interest. This lack of rigorous statistical methodology has limited the translational application and reproducibility of these methods.

Here, we present a statistical method and accompanying R package, `treeSeg`, that given a significance level *α*, tests for dependence of the response measurement distribution on all levels of hierarchy in a given tree while accounting for multiple testing. It returns the most likely segmentation of the tree such that each segment has a distinct response distribution, while controlling the overall false positive error. This is achieved by embedding the tree segmentation problem into a change-point detection setting (8–13).

`treeSeg` does not require any assumptions on the generation process of the tree *T*. It treats *T* as given and fixed, testing the response of interest against the given tree structure. Every tree *T*, independent of how it was generated, induces some latent ordering of the samples. `treeSeg` tests whether for this particular ordering, the distribution of the independent observations *y*_*i*_ depends on their locations on the tree.

`treeSeg` is applicable to a wide-range of problems across many scientific disciplines where the association between a tree structure and a response variable is under investigation, such as phylogenetic studies, molecular epidemiology, metagenomics, gene expression studies, etc. (3, 5, 14–16). The only inputs needed are the tree structure *T* and the outcome of interest *y*_*i*_ for the leaves of the tree. We demonstrate the sensitivity and specificity of `treeSeg` using simulated data and its application to a cancer gene expression study (1).

## 1. Results

For ease of presentation we restrict to discrete binary response measurements *y*_*i*_ ∈ {0, 1}. However, the procedure is equally applicable to continuous and other observation types (see Methods). If there is no association between the tree *T* and the response measurement *y*_*i*_, then the observed responses *y*_*i*_ would be randomly distributed on the leaves of the tree, independent of the tree structure *T*. However, if the distribution of responses is associated with the tree structure, we may observe clades in the tree with distinct response distributions. The power to detect segments with distinct distributions depends on the size of the clade and the change in response probability, *p*_*i*_ = **P**(*Y*_*i*_ = 1) = 1 − **P**(*Y*_*i*_ = 0), which means one can only make statistical statements on the *minimum* number of clades with distinct distributions on the tree and not the *maximum*. Figure 1 illustrates our method and its output for a simulated dataset. The responses *y*_*i*_ are displayed on the leaves of the tree as black and gray lines. The tree *T* is made of three segments with distinct distributions over the responses indicated by dark gray, light gray, and white backgrounds. Given a confidence level 1 − *α* (e.g., 1 − *α* = 0.9, 0.95), the `treeSeg` procedure estimates the most likely segmentation of the tree into regions of common response distributions, such that the true number of segments is at least as high as the estimated number of segments with probability of 1 − *α*.

**Fig. 1.**
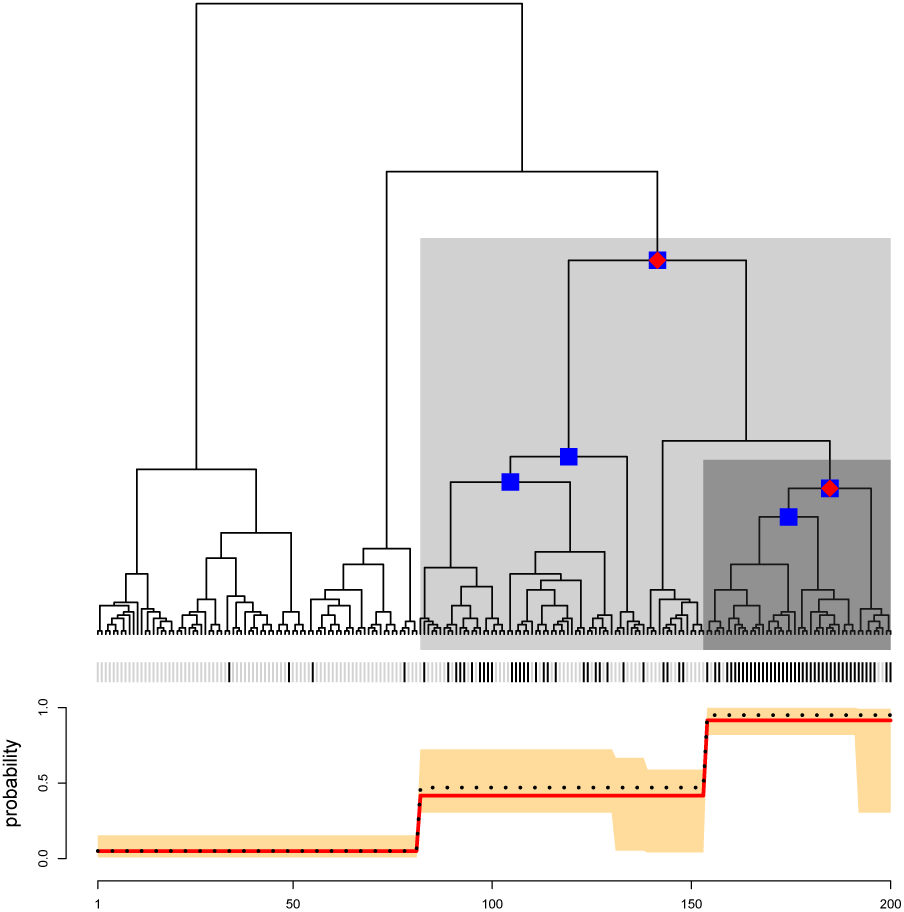
Illustration of the treeSeg method. Binary tree with 200 leaves and three segments with distinct response distributions indicated by dark gray, light gray and white backgrounds. Outcomes for each sample is shown on the leaves of tree as gray or black vertical lines. Leaf responses were simulated such that the black line has probability of 0.95, 0.47 and 0.05 for each of the dark gray, light gray and white background sections respectively. Using *α* = 0.1, `treeSeg` has estimated three segments on the tree with distinct response distributions indicated by the red dots on the nodes of the tree. Blue dots constitute a 90% confidence set for the nodes of the tree associated with the change in response distribution. The bottom panel shows the simulation response probabilities (black dotted line) and the `treeSeg` estimate (red line) and its 90% confidence bands in orange color.

Our method employs many likelihood ratio statistics simultaneously to test for changes in the response distribution on all levels of tree hierarchy and estimates at what level, if any, there is a change. The multiple testing procedure of `treeSeg` is based on a multiscale change-point methodology (11), tailored to the tree structure. The significance levels of the individual tests are chosen in such a way that the overall significance level is the pre-specified *α*. Theoretical proofs are provided in the Methods section and the online supplementary information. As well as the maximum likelihood estimate, our method also provides confidence sets (at 1 − *α* level) for the nodes of the tree associated with the change in response distribution and a confidence band for the response probabilities *p*_*i*_ over the segments.

In the example of Figure 1, using *α* = 0.1, `treeSeg` estimates three segments in the tree *T*, indicated with red dots on the nodes of the tree, recovering the true simulated changes in response distributions. In Figure 1, blue dots on the tree indicate the 1 − *α* confidence set for the nodes on the tree associated with the change in responses *y*_*i*_. The red line, in the bottom plot, shows the maximum likelihood estimate of the response probabilities *p*_*i*_ for each segment, which accurately recovers the true simulated probabilities shown as black dotted line. The orange band shows the 1 − *α* confidence band of the response probabilities.

The `treeSeg` method can handle missing data and to make response predictions on new samples. Computationally, `treeSeg` scales well with sample size, for example a test simulation for a tree with 100,000 samples (number of leaves in the tree) and no response association took around 110 minutes to run on a standard laptop. For details on `treeSeg`’s implementation see online supplementary text.

### A. Simulation Study

We confirmed the statistical calibration and robustness of `treeSeg` using simulation studies. We found that for reasonable minimal clade sizes and changes in response distribution, `treeSeg` is able to detect association between response and tree structure reliably (Supplementary Figures S11 - S16). More importantly, `treeSeg` almost never detects segments which are not present (it can be mathematically proven that the inclusion of a false positive segment only happens with probability ≤ *α*, see Methods) and the nominal guarantee of 1 − *α* is even exceeded in most cases (see online supplementary information). For example, in 1,000 randomly generated trees (200 leaves) with no changes in response distribution, `treeSeg` (using *α* = 0.1) correctly assigns no association between response and tree structure in 98.6% of the runs.

The `treeSeg` algorithm incorporates a fixed ordering of the leaf nodes according to the tree structure *T*. In principle, any such ordering is equally valid, as long as this is made a-priori, independent of the response variables. In our online implementation we apply a standardized pre-ordering of the nodes, so that `treeSeg`’s output is independent of any user specified ordering. Furthermore, we provide simulation results showing that `treeSeg` is very robust to random changes in node ordering and consistently infers the correct number of segments on the tree (Supplementary Figures S17 - S25). In Section E in the online supplement we also discuss an approach based on aggregating results among several random leaf orderings for the given tree. For more details on simulation studies refer to the online supplementary information.

### B. Application to Cancer data example

We illustrate the application of the method on a breast cancer gene expression study (1) where data are publicly available. Following the original study, we used correlation of gene expression data as a distance measure between samples to build a hierarchical clustering tree. In the original study, based on visual inspection, the authors divided the samples into two clusters, observing differences in the distributions of various clinical responses between the two clusters.

In contrast, `treeSeg` only requires a significance level *α* as input and searches for associations between responses and the tree on all levels of hierarchy, while accounting for multiple testing. Our results are shown in Figure 2. Using an *α* = 0.05, for one of the responses, `treeSeg` delinated the tree into two clusters with distinct response distribution as in the original study. However, `treeSeg` reports different patterns of association between the tree and the other five responses, including one, which has no association with tree structure.

**Fig. 2.**
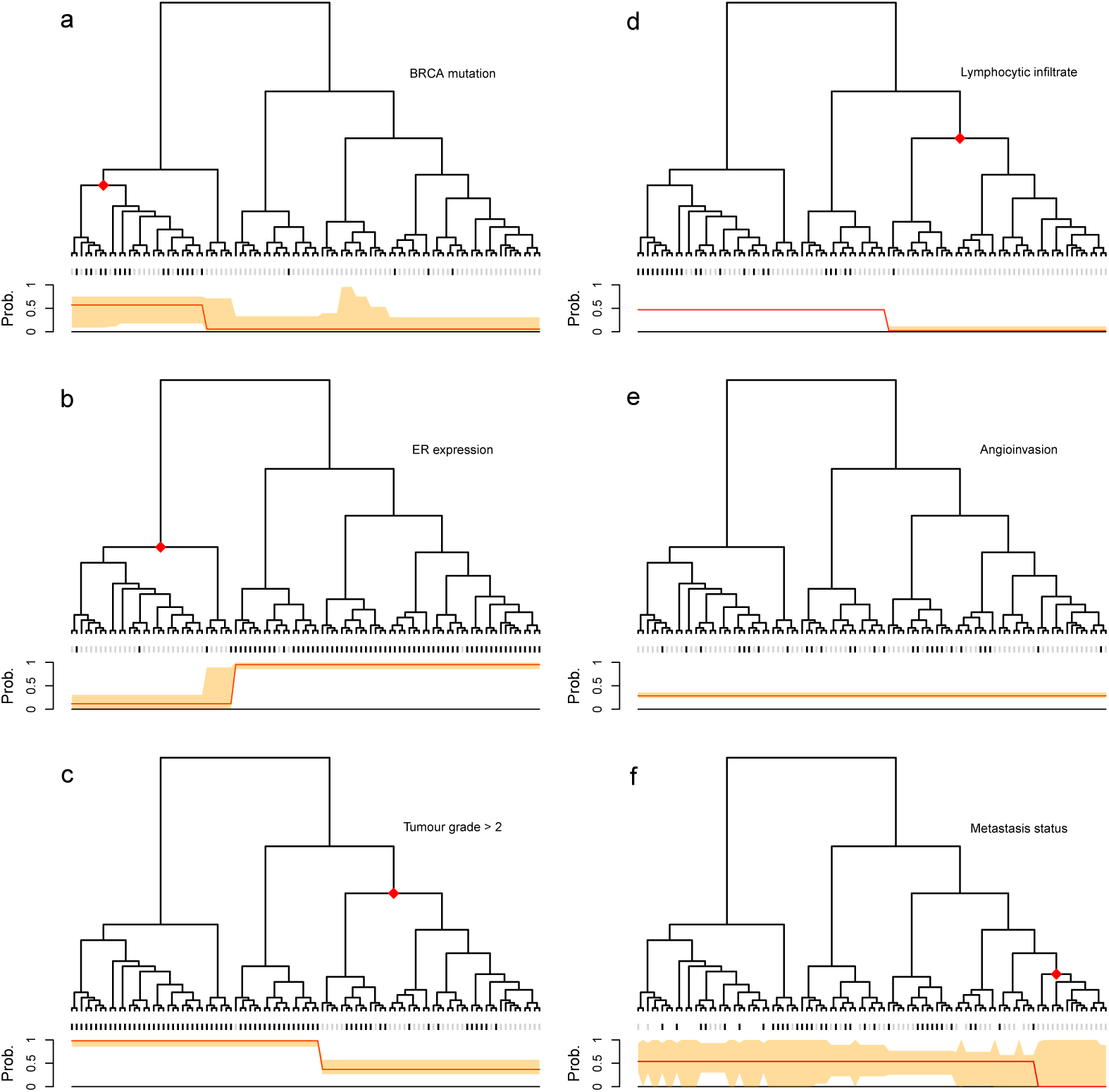
Application of `treeSeg` to a cancer gene expression study (1). Gene expressions for 98 breast cancer samples were clustered based on correlation between samples. Six clinical responses were collected for the samples, BRCA mutation, oestrogen receptor (ER) expression, histological grade, lymphocytic infiltration, angioinvasion and development of distant metastasis within 5 years which are represented in panels (a-f) respectively. In each panel, the `treeSeg` estimation (at *α* = 0.05) for clades with distinct response distribution and their probabilities are indicated by the red dots on the tree and the red line below the tree. The orange band shows the 95% confidence band for the response probabilities *p*_*i*_ for the estimated segments. In panel (e) there is no association between the tree and the response (angioinvasion) and in panel (f) some of the samples have missing observations for the response (distant metastasis within 5 years).

`treeSeg`’s application is not restricted to trees generated using hierarchical clustering. For an application to a maximum likelihood phylogenetic tree generated from pathogen sequence data see Section 3 in the online supplementary material. Indeed, `treeSeg` is completely independent of the particular way the tree was generated, with all results holding true conditioned on the specific tree *T* under consideration (see Section F in the online supplementary materials).

## 2. Discussion

The only tuning parameter for the `treeSeg` method is a significance level *α*. Depending on the application, the user can decide which value of *α* is appropriate, or screen through several values of *α*, e.g., *α* = 0.01, 0.05, 0.1, 0.5. A small *α* gives a higher confidence that all detected associations are, indeed, present in the data (with probability of at least 1 − *α*). A larger *α* allows to detect more clusters, but increases the risk of including false positive clusters.

The confidence statement for detected clades and response probabilities that accompany `treeSeg`’s segmentation, account for multiple testing at the level of 1 − *α*. This allows for precise uncertainty quantification when detecting associations between tree structure and the responses. We highlighted `treeSeg`’s potential with an example from a gene expression study, but note its ubiquitous applicability in various settings. Our method `treeSeg` is implemented as an R package, available on GitHub (https://github.com/merlebehr/treeSeg) and accompanied by a detailed Jupyter notebook with reproductions of all figures in this text, and has the potential to be used across many fields of science.

## 3. Methods

### A. Model assumptions

For illustration purposes, we focus on binary traits *Y*_*i*_ ∈ {0, 1}. Please see online supplementary text Section B on how `treeSeg` generalizes for arbitrary continuous and discrete data. We assume a fixed given rooted tree *T* with *n* leaves, that captures some neighborhood structure of interest. For the *n* samples (the leaves of the tree) independent binary traits *Y*_*i*_, *i* = 1,…, *n*, with success probability *p*_*i*_, are observed, that is,

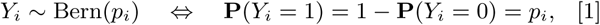

independently for *i* = 1,…, *n*, where Bern(*p*) denotes a Bernoulli distribution with success probability *p*. The aim is to estimate the underlying success probabilities *p*_1_,…, *p*_*n*_ from the observations *Y*_*i*_. Without any additional structural information on the success probabilities we cannot do better than estimating *p*_*i*_ = *Y*_*i*_. However, taking into account the tree structure, we can assume that the success probabilities are associated with the tree such that samples on the same clade of the tree may have the same success probabilities. Our methodology is based on a testing problem, where the null model assumes that all isolates have the same success probability, say *p*_0_, and the alternative model assumes that some of the clades on the tree have different success probabilities (*p*_0_ + *c* ∈ [0, 1]). In the following, we denote an internal node (which demarcates a clade on the tree) with a distinct success probability as an *active node*.

For simplicity, we will assume in the following that the tree *T* is binary. Extensions to arbitrary trees are straight forward. We use the following notation. For a binary, rooted tree *T* = (*V, E*) we assume vertices *V* = 1,…, *n, n*+1,…, 2*n*−1 and edges *E* = (*i, j*) : *i, j* ∈ *V* with *i, j* connected}. The leaves are labeled *V*_*L*_ = {1,…, *n*}, the inner nodes are labeled *V*_*I*_ = {*n* + 1,…, 2*n*−1}, and the root is labeled 2*n*−1. For a node *i* ∈ *V* its set of offspring leaves in *V*_*L*_ is denoted as Off(*i*). For a node *i* ∈ *V* the subtree of *T* with root *i* is denoted as *T* (*i*). An illustrative example for this notation is shown in Figure 3. Moreover, for an inner node *i* ∈ *V*_*I*_ with offspring leaves Off(*i*) = {*i*_1_,…, *i*_*m*_} ⊂ {1,…, *n*} and for some *ϵ* ∈ (0, 1), we denote the left *ϵ*-leaf-neighborhood of *i* as N_L_(*i, ϵ*) = {*i*_1_−⌊*nϵ*⌋, *i*_1_−⌊*nϵ*⌋ + 1,…, *i*_1_ + ⌊*nϵ*⌋−1, *i*_1_ + ⌊*nϵ*⌋} and, analog, the right *ϵ*-leaf-neighborhood of *i* as N_R_(*i, ϵ*) = {*i*_*m*_ −⌊*nϵ*⌋, *i*_*m*_−⌊*nϵ*⌋ + 1,…, *i*_*m*_ + ⌊*nϵ*⌋−1, *i*_*m*_ + ⌊*nϵ*⌋}.

We consider the following statistical model.

**Fig. 3.**
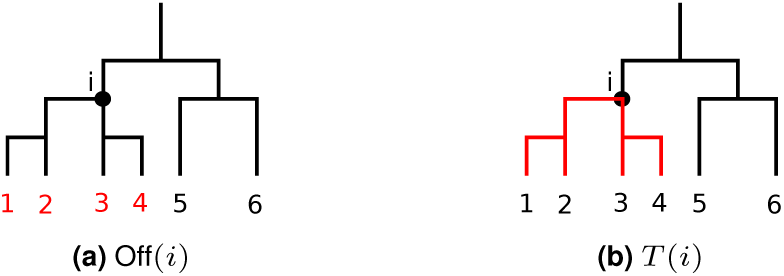
Illustration of notation used in the paper. The respective subset of leaf nodes Off(*i*) and sub-tree *T* (*i*) are shown in red.

#### Model 1

For a given binary, rooted tree *T* = (*V, E*) as above, assume one observes for each of the leaves *i* ∈ *V*_*L*_ independent Bernoulli random variables *Y*_*i*_ ∼ *B*(*p*_*i*_), 1 ≤ *i* ≤ *n*, where the vector of success probabilities *p* = (*p*_1_,…, *p*_*n*_) is an element of

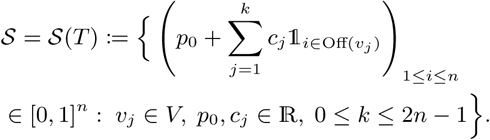

For an element *p* ∈ 𝒮 we denote the set of nodes *V* (*p*) := {*v*_1_,…, *v*_*k*_} as a set of *active nodes* and *k*(*p*) = *k* as the number of active nodes. To ensure identifiability of active nodes, we further assume that for each active node *v*_*j*_, *j* = 1,…, *k*, there exists at least one offspring leaf *i* ∈ 1,…, *n* that has the same success probability as *v*_*j*_. This just excludes the trivial case where the influence of one active node (or the root) is completely masked by other active nodes. Equivalently, this means that we assume #supp(*p*) = # {*p*_*i*_ : *i* = 1,…, *n*} = *k* + 1. We provide a simple example in the supplementary material Figure S27.

We stress that the set *V* (*p*) is not necessarily unique (see Figure S26 in the supplement for an example). That is, for a given vector *p* ∈ 𝒮 there may exists two (or more) sets of active nodes {*v*_1_,…, *v*_*k*_} and 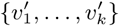 such that

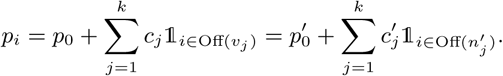

To overcome this ambiguity we will implicitly associate with each *p* ∈ 𝒮(*T*) a set of active nodes *V* (*p*) of size *k*(*p*). When a specific vector *p* ∈ 𝒮(*T*) has more than one possible sets of active nodes *V* ′, *V* ″,…we assume several copies of *p* in 𝒮(*T*), one associated with each of the sets *V* ′, *V* ″, In the following, we explore the tree structure to estimate the underlying success probabilities *p*_*i*_ and hence, their segmentation into groups of leaves where observations within a group have the same success probability and observations between different groups have different success probabilities.

### B. Multiscale segmentation

The procedure we propose extends (11) from totally ordered structures to trees and is a hybrid method of estimating and testing. A fundamental observation is that one can never rule out an additional active node. This is because a node could be active but changed the success probability of its offspring nodes only by an arbitrarily small amount. On the other hand, if in a subtree successes are much more common than in the remaining tree, it is possible to significantly reject the hypothesis that all leaves in the tree have the same success probability.

For a given candidate vector 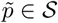 our procedure employs on each subtree *T*(*i*) where 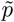 is constant, with 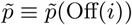, a likelihood ratio (LR) test for the hypothesis that the corresponding observations all have the same success probability 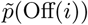. The levels of the individual tests are chosen in such a way that the overall level of the multiple test is *α*, for a given pre-specified *α* ∈ (0, 1). A statistical hypothesis test can always be inverted into a confidence statement and vice versa. Therefore, we can derive from the above procedure a confidence set for the vector of success probabilities *p* = (*p*_1_,…, *p*_*n*_). We require our final estimate 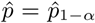 to lie in this confidence set. That is, we require whenever 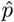 has constant success probability on a subtree, the respective LR test accepts. Within all vectors *p* ∈ 𝒮 which lie in this confidence set we choose one which comes from a minimum number of active nodes and within this set, we choose the maximum likelihood solution. Thereby, our procedure does not only provide an estimate, but also confidence statement for all quantities (11). More precisely, the following asymptotic confidence statements hold true.

1. With probability at least 1 − *α* the true underlying signal *p* ∈ 𝒮 originates from at least 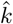 active nodes, where 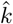 is the number of active nodes of 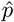, see Theorem 1.
2. `treeSeg` yields a set of nodes 𝒞_1−α_, such that the active nodes of *p, V* (*p*), are contained with probability at least 1 − *α* in 𝒞_1−α_, see Corollary 4.
3. `treeSeg` yields a confidence band for the underlying signal *p*, denoted as <u>*p*</u>_1−*α*_ and 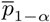, such that with probability at least 1 − *α* it holds that 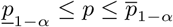 simultaneously for all *i* = 1,…, *n*, see Theorem 3.

Moreover, the coverage probability of the confidence sets allows to derive (up to log factors) optimal convergence rates of the `treeSeg` estimator as the sample size *n* increases (11). In particular, we show the following.

1. For fixed overestimation bound *α* ∈ (0, 1), the probability that `treeSeg` underestimates the number of active nodes *k* vanishes exponentially as *n* increases, see Theorem 2.
2. The localization of the estimated active nodes is optimal up to a leaf node set of order log(*n*), see Theorem 5.

In the following we will give details of the method and of the statements 1–5. The proofs of Theorem 1, 2, 3, 5 and Corollary 4 are similar to the totally structured setting (11). Necessary modifications are outlined in the online supplementary text.

For an arbitrary given test vector 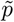 (which may depend on *Y*) we define the multiscale statistic (11, 17–19)

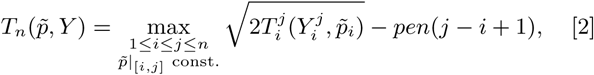

where 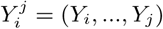 and 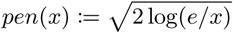. Here, 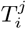 is the local log-LR test statistic (20) for the testing problem

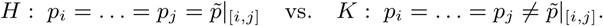

The calibration term *pen*(·) serves as a balancing of the different scales in a way that the maximum in [2] is equally likely attained on all scales (11, 17) and guarantees certain optimality properties of the statistic [2] (17). Assuming a minimal segment scale *λ* ∈ (0, 1) of the underlying success probability vector *p*, that is,

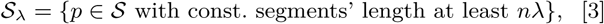

it can be shown that *T*_*n*_(*p, Y*) converges in distribution to a functional of the Brownian motion (11) which is stochastically bounded by

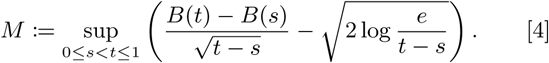

Thereby, the minimal scale *λ* may depend on *n* such that *nλ/log*(*n*)^3^ → ∞ as *n* → ∞, see (11). As the distribution of *M* does not depend on the true underlying signal *p*, its quantiles can be obtained by Monte Carlo simulations and are, in the following denoted as *q*_1−α_, that is,

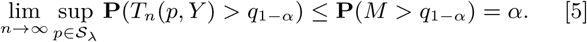

For a given confidence level *α* ∈ (0, 1) or, equivalently, a threshold value *q* = *q*_1−*α*_ in [5], we first define an estimator for the number of active nodes *k* in Model 1 via

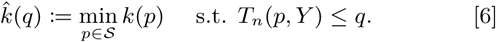

Once the number of active nodes *k* is estimated, we estimate *p* as the constrained maximum likelihood estimator

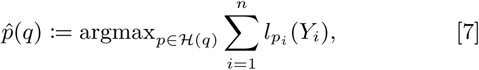

where *l*_*p*_(*y*) is the log-likelihood function of the binomial distribution and

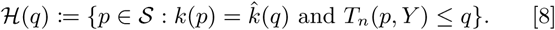

Note that the maximum likelihood (ML) solution in [7] is not necessarily unique. On the one hand, this is due to the non-uniqueness of the active nodes. On the other hand, this might happen with positive probability by the discreteness of the Bernoulli observations *Y*. In that case, `treeSeg` just reports the first available solution, with all other equivalent solutions listed in the confidence set. Clearly, if we choose *q* as in [5] for some given confidence level *α* ∈ (0, 1) the estimator 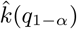 asymptotically controls the probability to overestimate the number of active nodes, as summarized in the following theorem.

#### Theorem 1.

For fixed minimal scale *λ* > 0 and significance level 1 − *α* ∈ (0, 1), let 𝒮_*λ*_ be as in [3], *q*_1−*α*_ as in [5], and 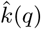 `treeSeg`’s estimated number of active nodes in [6]. Then it holds that

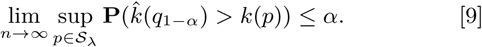

We stress that in Theorem 1 it is possible to let *λ* go to zero as *n* increases (11), recall the paragraph after equation [4]. In particular, from the construction of 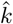 in [6] it follows that *T*_*n*_(*p, Y*) ≤ *q* implies 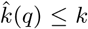 and thus, for the set ℋ(*q*_1−*α*_) in [8] one obtains that

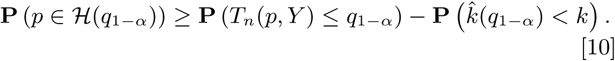

By [5] it follows that the first term on the right hand side, **P** (*T*_*n*_(*p, Y*) ≤ *q*_1−*α*_), is asymptotically lower bounded by 1 − *α*. Moreover, as we show in the following Theorem 2, the underestimation error 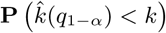 vanishes exponentially fast as sample size *n* increases. From this it follows that the set ℋ(*q*_1−*α*_) constituents an asymptotically honest confidence set (11) for the whole vector *p* from which confidence sets as in statements 2 and 3 follow, see Theorem 3 and Corollary 4.

Any bound on the underestimation error necessarily must depend on the minimal segment scale *λ* in [3], as well as a minimal pairwise difference *δ* ∈ (0, 1) of success probabilities in different active segments. That is, for *p* ∈ 𝒮_*δ*_ we assume 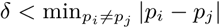 and let

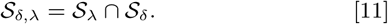

With this, one obtains that the underestimation probability decreases exponentially (11) in *n* (for fixed *δ, λ* and significance level 1 − *α* ∈ (0, 1)) as the following theorem shows.

#### Theorem 2.

For fixed minimal scale *λ* > 0, minimal success probability difference *δ* > 0, and significance level 1 − *α* ∈ (0, 1), let 𝒮_*λ,δ*_ be as in [11], *q*_1 − *α*_ as in [5], and 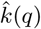 `treeSeg`’s estimated number of active nodes in [6]. Then it holds that

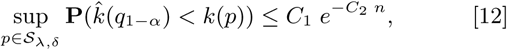

where *C*_1_ and *C*_2_ are positive constants, which only depend on *α, λ, δ*.

Again, it is possible to let *α, λ*, and *δ* go to zero as the sample size *n* increases (11). The proof of Theorem 2 is similar as for totally ordered structures (11). We outline necessary modifications in the online supplementary text. From Theorem 2 and [10] we directly obtain that ℋ(*q*_1−*α*_), indeed, constitutes a 1 − *α* asymptotic confidence set for the segmentation *p*.

#### Theorem 3.

For fixed minimal scale *λ* > 0, minimal success probability difference *δ* > 0, and significance level 1 *α* (0, 1), let 𝒮_*λ,δ*_ be as in [11], *q*_1−*α*_ as in [5], and (*q*_1−*α*_) as in [8], then

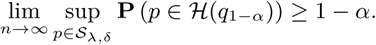

As a corollary, we also obtain a confidence set of the active nodes.

#### Corollary 4.

For fixed minimal scale *λ* > 0, minimal success probability difference *δ* > 0, and significance level 1− ∈ *α* (0, 1), let 𝒮_*λ,δ*_ be as in [11], *q*_1−*α*_ as in [5], and ℋ(*q*_1−*α*_) as in [8], then

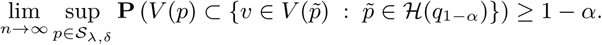

Theorem 1 and 2 reveal `treeSeg`’s ability to accurately estimate the number of active nodes in Model 1. For any (arbitrarily small) *α* ∈ (0, 1) we can control the overestimation probability by 1 − *α*, see Theorem 1. Simultaneously, as the sample size *n* increases, the underestimation error probability vanishes exponentially fast, see Theorem 2. The next theorem shows that `treeSeg` does not just estimate the number of active nodes correctly with high probability, but that it also estimates the location of those active nodes with high accuracy. To this end, note that for any active node *v* ∈ *V* (*p*) and any *ϵ* ≥ 1*/n* the leaf nodes of its left *ϵ*-leaf-neighborhood N_L_(*v, ϵ*) have non-constant success probability. The same is true for the right *ϵ*-leaf-neighborhood N_R_(*v, ϵ*). Now assume that `treeSeg` estimates the number of active nodes correctly 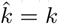, which is the case with high probability by Theorem 1 and 2. Then, 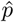 being non-constant on both, N_R_(*v*, 1*/n*) and N_L_(*v*, 1*/n*), for any true active nodes *v* ∈ *V* (*p*), implies a perfect segmentation. Thus, the following theorem shows that `treeSeg`, indeed, yields such a perfect segmentation up to a leaf node set of size 𝒪(log(*n*)). That is, conditioned on the correct model dimension 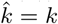, `treeSeg`’s segmentation is perfect up to at most an order of log(*n*) miss-classified leaf nodes.

#### Theorem 5.

For fixed minimal scale *λ* > 0, minimal success probability difference *δ* > 0, and significance level 1−*α* ∈ (0, 1), for any *p* ∈ 𝒮_*λ,δ*_ it holds true that

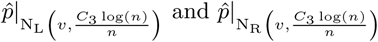

are not constant, for all *v ∈ V* (*p*), with probability at least 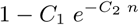, where *C*_1_, *C*_2_, *C*_3_ are positive constants which only depend on *α, λ, δ*.

Theorem 5 follows directly by translating change-point location estimation accuracy results for the totally ordered case (11) to the tree setting.

A natural question is whether the localization rate in Theorem 5 is optimal. In particular, one can compare this result to the totally ordered setting, where the minimax optimal change-point estimation rate is known to be of the same order (possibly up to log(*n*)-factors). One would expect that the additional tree structure leads to a strictly better segmentation rate. It turns out, however, that, without making further assumptions on the particular tree, the rate in Theorem 5 cannot be improved in general, see Theorem 6 in the supplement. On the other hand, when one imposes additional structural assumptions on the tree, for example, for perfect trees, it can be shown that `treeSeg` yields a perfect segmentation with high probability, see Theorem 7 in the supplement. Thus, `treeSeg` efficiently leverages the tree structure to overcome the minimax lower bound from a simple change-point estimation problem, whenever the tree allows this. We provide more details in Section D in the supplement.

## ACKNOWLEDGMENTS

MB was supported by DFG postdoctoral fellowship BE 6805/1-1. Moreover, MB acknowledges funding of DFG-GRK 2088. AM was funded by the Deutsche Forschungsgemeinschaft (DFG, German Research Foundation) under Germany’s Excellence Strategy - EXC 2067/1-390729940. AM and MB acknowledge support of DFG-SFB 803 Z02. The authors thank Laura Jula Vanegas for help with parts of the implementation.

