## Supplemental Text for "Testing for dependence on tree structures"

#### This PDF file includes:

Supplementary text
Figs. S1 to S27
References for SI reference citations

### Supporting Information Text

### 1. Methods

For the following, we need two additional notations. For an inner node $i \in V_I$ the two (direct) offspring nodes $i_1, i_2$ are denoted as $\mathrm{off}(i)$. For an inner node $i \in V_I$ the nodes of $T(i)$, recall Figure 3, without $i$ are denoted by $\mathrm{OFF}(i)$. An illustrative example for this notation is shown in Figure S1.

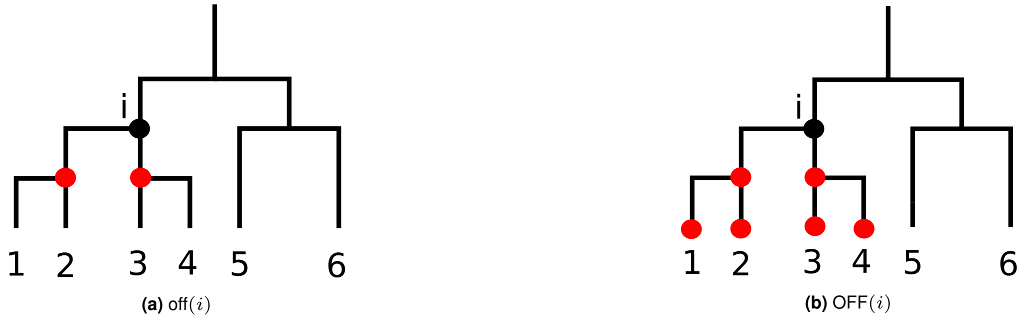


**Fig. S1.** Illustration of notation. The respective subset of nodes denoted by off($i$) and OFF($i$) are shown in red.

**A. Implementation.** The proposed multiscale procedure can be computed with dynamic programming (DP). The key idea of DP is to make use of overlapping subproblems. Here, the canonical related subproblems $P_m$ for $m \in V_I$ only consider those observations $Y_j$ where leaf $j$ is an offspring of inner node $m$, that is, $j \in \mathrm{Off}(m)$ . Let $\hat{p}^m$ denote the `treeSeg` solution for subproblem $P_m$. The problems $P_m$ are partially ordered. For an inner node $m$ with problem $P_m$ the subordinate problems $P_i$ are those where $i$ is an offspring of $m$ in the tree, that is $i \in \mathrm{OFF}(m)$. The relation between these problems $P_m$ (usually denoted as Bellman's equation in DP) is as follows: For a given inner node $m$ the solutions of $P_l$, $l \in \mathrm{OFF}(m)$ relate to the solution of $P_m$ via

$$\hat{p}^m \in \left\{ \left( c + \sum_{j=1}^{k} \hat{p}_i^{l_j} \mathbb{1}_{\{i \in \mathrm{Off}(l_j)\}} \right)_{1 \le i \le n} \middle| \; l_j \in \mathrm{OFF}(m),\; l_j \notin \mathrm{Off}(l_{j'}) \forall j \neq j',\; c \in [0,1] \right\}.$$

This means that if $l \in \mathrm{OFF}(m)$ is an active node of $\hat{p}^m$, the solution $\hat{p}^l$ of problem $P_l$ equals $\hat{p}^m$ on the offspring leaves of $l$. Thus, solving $P_m$ only amounts to finding the position(s) of the last active node(s). Here, an active node is *last* if there is no other active node which is its ancestor. Employing this DP scheme, the estimator $\hat{p}$ in (7) can be computed by successively solving the subproblems $P_m$, starting at the leaves and ending at the root $m = 2n - 1$ with final solution $\hat{p} = \hat{p}^{2n-1}$. In that way, one makes use of overlapping subproblems without having to run exhaustively over all possible $\mathcal{O}(n^n)$ tree segmentations. For each of the $\mathcal{O}(n)$ subproblems $P_m$, inner node $m$ has at most $\mathcal{O}(n)$ offspring nodes. Thus, in the worst case, solving $P_m$ amounts to consider $\mathcal{O}(n^{\hat{k}})$ *last* active nodes, where for each the multiscale constraint can be checked in $\mathcal{O}(n)$ time (1). To this end, note that $\hat{k}(m) \in \{\hat{k}(i_1) + \hat{k}(i_2), \hat{k}(i_1) + \hat{k}(i_2) + 1\}$ where $\{i_1, i_2\} = \mathrm{off}(m)$ are the two direct offspring nodes of $m$ and $\hat{k}(m)$ denotes the estimated number of active nodes in subproblem $P_m$. In total, this gives a worst case run time of $\mathcal{O}(n^{\hat{k}+2})$. However, by keeping track of the individual confidence sets $\mathcal{C}_{1-\alpha}^m$ of problem $P_m$, in practice, only a small fraction of the $n^{\hat{k}}$ possible segmentations has to be considered. This renders the actual runtime to be usually much faster and makes this DP scheme very efficient in practice.

**B. Extensions for continuous and other discrete distributions.** The `treeSeg` methodology is not restricted to binary response variables and can be applied to various other types of continuous and discrete distributions. When the response distribution belongs to a one-parametric distributional family, that is, one observes independently $Y_i \sim F_{\theta_i}$, where $F_\theta$ belongs to a distributional family with parameter $\theta$, `treeSeg` can segment with respect to $\theta$ by simply replacing the log-LR tests $T_i^j$ in (2) by the one for the testing problem

$$H : \theta_i = \ldots = \theta_j = \tilde{\theta}|_{[i,j]} \quad \text{vs.} \quad K : \theta_i = \ldots = \theta_j \neq \tilde{\theta}|_{[i,j]}.$$

The respective multiscale statistic $T_n$ in (2) still converges in distribution with a limit bounded by $M$ in (4) under very weak assumptions (in particular, whenever $F_\theta$ belongs to an exponential family) and hence, all confidence statement remain valid (1, 2). In the current `treeSeg` R package an implementation for continuous traits under a Gaussian distribution assumptions is available. Figure S2 shows a similar example as in the main text, but with normally distributed response. We are planning to add implementations for various other distributional families to the `treeSeg` R package in the future, such as exponential and Poisson. Moreover, recently, a multiscale procedure for quantile segmentation of arbitrary independent traits was proposed (3) and we are also planning to implement this in the future for `treeSeg`.

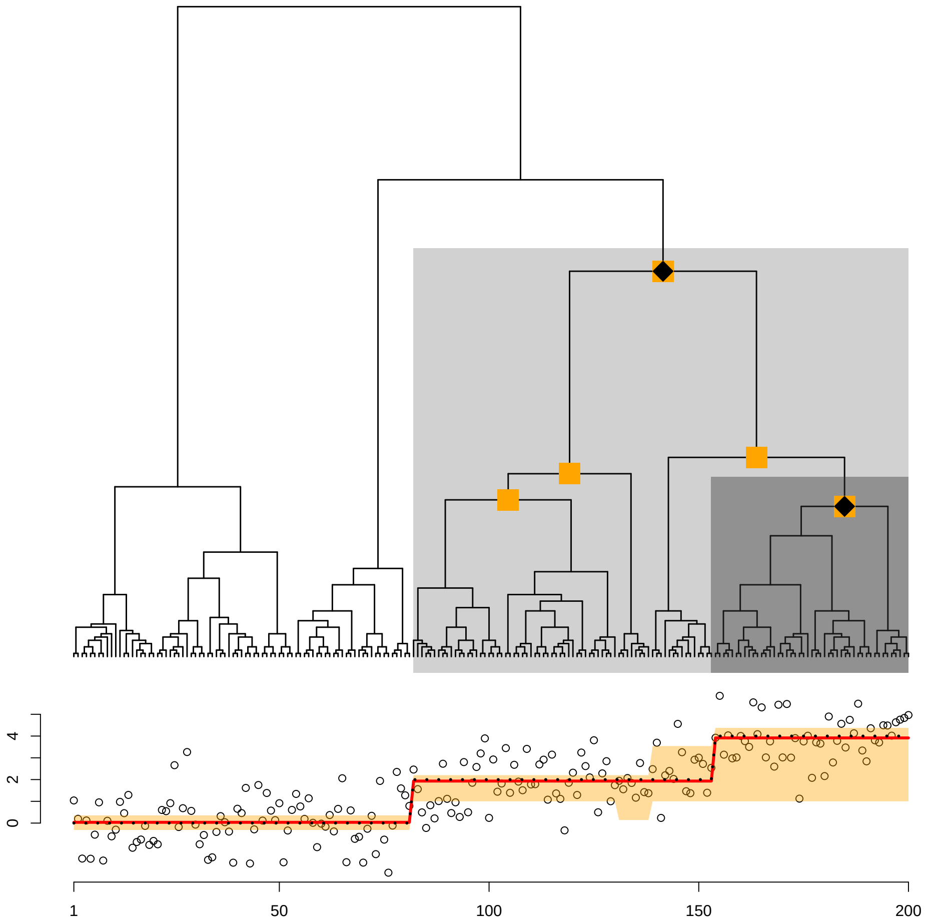


**Fig. S2.** Illustration of the `treeSeg` method for continuous data analog as in example of the main text. Leaf phenotype samples (black dots in bottom panel) come from a normal distribution with variance 1 and mean equal to the black dotted curve. Using $\alpha = 0.1$, `treeSeg` estimates the correct three segments on the tree with distinct phenotype distributions indicated by the black dots on the nodes of the tree. Orange nodes constitute a 0.90% confidence set for the nodes of the tree associated with the change in phenotype distribution. The bottom panel shows the `treeSeg` estimate for phenotype mean (red line) and its 90% confidence band (orange area).

**C. Proof of Theorem 2.** In the following we outline necessary modifications of the proof of Theorem 2 for tree structured signals $p \in \mathcal{S}$ compared to totally ordered signals (1). In the change-point detection problem for totally ordered structures (1) the number of active nodes relates to the number of change-points of the underlying signal. There, one shows the underestimation bound as in Theorem 2 as follows: Consider a true underling signal with $k$ change-points and each of the $k+1$ constant segments having at least length $\lambda n$. For each of the $k$ change-points consider an interval $\mathcal{I}_i$, $i = 1, \ldots, k$, of length $\lambda n$ centered around this change-point. Note that those $k$ intervals will be disjoint and on each of them the true underlying signal makes a jump of size at least $\delta$. Now, when an estimated segmentation has less then $k$ change-point, this implies that for at least one of those $k$ intervals $\mathcal{I}_i$, the estimator is constant taking some value $c \in \mathbb{R}$. This implies that the two multiscale tests on the left and on the right half of $\mathcal{I}_i$, with true signal-difference of at least $\delta$, both accepted the constant value $c$. It follows from power considerations of the LR test that this event has probability vanishing exponentially in $n$ (for fixed $\lambda, \delta, \alpha$).

In our setting, $k$ corresponds to the number of active nodes, which, in general, can be different to the number of change-points in the leaf-signal $(p_i)$, $i = 1, \ldots, n$. Note that, depending on the location of the active node $v \in V(p)$, it may induce either one or two change-points in $p$. In order to apply the same proof as in the totally order case (1), for a fixed underlying signal $p \in \mathcal{S}$, we have to find intervals $\mathcal{I}_1, \ldots, \mathcal{I}_m$, such that

a) $p$ is non-constant on $\mathcal{I}_i$,

b) $p$'s constant segments in $\mathcal{I}_i$ have minimal length $C\lambda n$ for some constant $C$,

c) any other $\tilde{p} \in \mathcal{S}$ with less then $k$ active nodes, that is $k(\tilde{p}) < k$, will be constant on at least one of the intervals $\mathcal{I}_1, \ldots, \mathcal{I}_m$.

For $p \in \mathcal{S}$ one can construct regions $\mathcal{I}_1, \ldots, \mathcal{I}_m$ which fulfill a)-c) as follows. To this end, formally write the root $v_0 = 2n + 1$ as an additional active node. Further, let $C$ be some sufficiently small constant (which may depend on $\lambda$).

1. For each change-point of $(p_1, \ldots, p_n)$ select an interval $\mathcal{I}_i \subset \{1, \ldots, n\}$ of length $C\lambda n$ centered at the change-point.

2. Moreover, for each pair of true active nodes $v_i \neq v_j \in V(p)$, $i, j = 0, \ldots, k$ select a union of two intervals $\mathcal{I}_{v_i,v_j} = \mathcal{I}^{v_i}_{v_i,v_j} \cup \mathcal{I}^{v_j}_{v_i,v_j} \subset \{1, \ldots, n\}$, with $\mathcal{I}^{v_i}_{v_i,v_j}$ a constant region from active node $v_i$ of minimal length $C\lambda n/2$ and $\mathcal{I}^{v_j}_{v_i,v_j}$ accordingly.

It is easy to see that these intervals satisfy a) and b). In the following we show by induction that these intervals also satisfy property c).

The base case, for $k = 1$, follows trivially. Now, as induction hypothesis, assume c) holds true for all $k = 1, \ldots, \kappa - 1$. Let $v_1, \ldots, v_\kappa$ be the true active nodes of some underlying signal $p \in \mathcal{S}$. Assume there exists some $\tilde{p}$ with active nodes $\tilde{v}_1, \ldots, \tilde{v}_{\kappa-1}$ such that $\tilde{p}$ is not constant for all intervals $\mathcal{I}_i$, $\mathcal{I}_{v_i,v_j}$. We denote the *influence region* IR of true (and estimated, respectively) active node $v_i$ (and $\tilde{v}_i$, respectively) as those leaf nodes, which are a direct offspring of $v_i$ (and $\tilde{v}_i$, respectively), that is, which have the same success probability as $v_i$ (and $\tilde{v}_i$, respectively). See Figure S4 for an example. We say that estimated active node $\tilde{v}_i$ is *related to* true active node $v_i$ if the influence regions of $\tilde{v}_i$ and $v_i$ intersect, that is $\mathrm{IR}(\tilde{v}_i) \cap \mathrm{IR}(v_i) \neq \emptyset$. See Figure S3 for an example. That is, when $v_i$ is either a direct offspring of $\tilde{v}_i$ (without other estimated active nodes in between) or when $\tilde{v}_i$ is a direct offspring of $v_i$ (without other true active nodes in between).

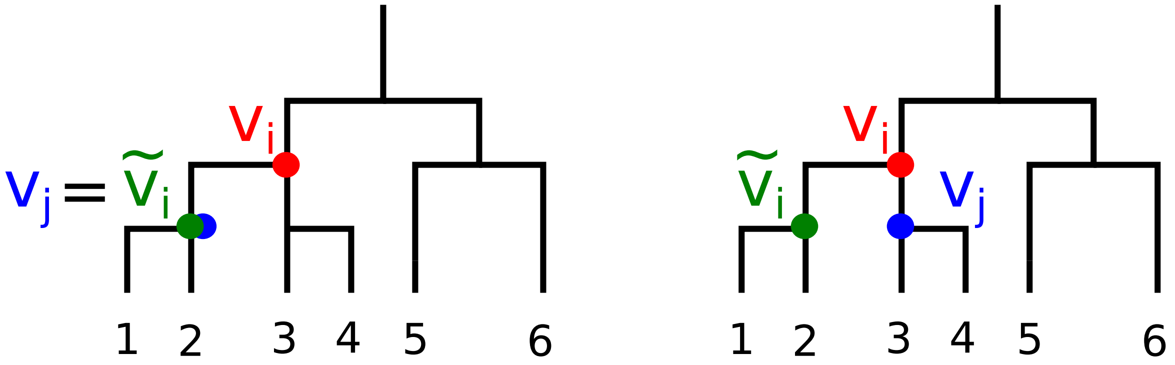


**Fig. S3.** Example of tree with two true active nodes $v_i, v_j$ and one estimated active node $\tilde{v}_i$. In the right example $v_i$ and $\tilde{v}_i$ are related. In the left example $v_i$ and $\tilde{v}_i$ are not related.

Assume that there exists one estimated active node $\tilde{v}_i$ which is related to exactly one true active node $v_j$, $j = 1, \ldots, \kappa$. Then, for the reduced problem with true $p' \in \mathcal{S}_{\delta,\lambda}$ having active nodes $\{v_1, \ldots, v_\kappa\} \setminus \{v_j\}$, there exists $\tilde{p}'$ with active nodes $\{\tilde{v}_1, \ldots, \tilde{v}_{\kappa-1}\} \setminus \{\tilde{v}_i\}$, such that $\tilde{p}'$ is non-constant on all the intervals $\mathcal{I}_1, \ldots, \mathcal{I}_{m'}$ of $p'$. This is a contraction to the induction hypothesis. The same contraction follows when there exists one estimated active node $\tilde{v}_i$ which is related to no true active node $v_j$. Consequently we deduce the following.

- *Every estimated active node $\tilde{v}_j$ is related to two true active nodes $v_{j_1}, v_{j_2}$, $j_1, j_2 \in \{1, \ldots, \kappa\}$.*

Now, consider some estimated active node $\tilde{v}_j$ which has no other offspring estimated active nodes. Let $v_{j_1}$ and $v_{j_2}$ be two related true active nodes of $\tilde{v}_j$. If $v_{j_1}$ and $v_{j_2}$ are both offspring of $\tilde{v}_j$, it follows that $\tilde{p}$ is constant on $\mathcal{I}_i$ which connects $v_{j_1}$ and $v_{j_2}$, which is a contraction. If $v_{j_1}$ and $v_{j_2}$ are both ancestors of $\tilde{v}_j$, it follows that $\tilde{v}_j$ is not related with one of $v_{j_1}$ and $v_{j_2}$, which is also a contradiction. Consequently, without loss of generality, one can assume that $v_{j_1}$ is an ancestor of $\tilde{v}_j$ and $v_{j_2}$ is an offspring of $\tilde{v}_j$, with $j_1, j_2 = 1, \ldots, \kappa$. One can applying the same argument successively for all estimated active nodes starting at the leaves and ending at the root to obtain the following.

- *All estimated activated nodes have both, an offspring and an ancestor true active node.*

In particular, it also follows that $\tilde{v}_j$ cannot be related with the root active node $v_0$ (as it is an offspring of $v_{j_1}$), which yields the following.

- *Every true active node $v_i$ is related with at least one estimated active node $\tilde{v}_j$.*

To see this note that otherwise $\tilde{p}$ would be constant on the regions which connects $v_i$ and $v_0$. Now, consider some true active node $v_{i_1}$ which has no other offspring true active nodes. Let $\tilde{v}_i$ be its related estimated active node. Note that $\tilde{v}_i$ has to be an ancestor of $v_{i_1}$ as otherwise $\tilde{v}_i$ would have no offspring true active node. Further, let $v_{i_2}$ be an ancestor true active node of $\tilde{v}_i$. As $\tilde{p}$ is non-constant on $\mathcal{I}_i$ which connects $v_{i_1}$ and $v_{i_2}$, it follows for the reduced problem with true signal $p' \in \mathcal{S}_{\delta,\lambda}$ having active nodes $\{v_1, \ldots, v_\kappa\} \setminus \{v_{i_1}\}$, that there exist $\tilde{p}'$ with active nodes $\{\tilde{v}_1, \ldots, \tilde{v}_{\kappa-1}\} \setminus \{\tilde{v}_i\}$ which is non-constant on all the intervals $\mathcal{I}_1, \ldots, \mathcal{I}_{m'}$ of $p'$. This is a contraction to the induction hypothesis and finishes the induction step.

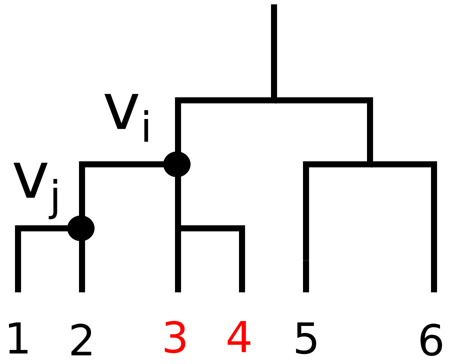


**Fig. S4.** Example of tree with two active nodes $v_i, v_j$. The *influence region* of $v_i$, IR($v_i$), is marked red.

**D. Comparison of estimation rates for tree structures vs. totally ordered change-point structures.** The `treeSeg` procedure incorporates into its analysis a specific ordering of the leaf nodes $i = 1, \ldots, n$ that is induced by the tree $T$. If one was only interested in the change-points of the leaf nodes that correspond to this ordering [*], one could also just apply a standard change-point procedure to the ordered observations $Y_1, \ldots, Y_n$, without taking any further information of the tree $T$ into account.

A natural question is how the statistical change-point estimation rates of `treeSeg` compare to those of a simple change-point procedure. In particular, one would hope that the additional information from the tree $T$ (beyond just such a simple ordering of the leaf nodes), improves the statistical estimation accuracy for the segmentation. In the following we provide two results which give insight into this question:

- First, we show that without making any further assumptions on the tree $T$, the statistical segmentation rate of order $1/n$ that we show for `treeSeg` in Theorem 5 up to the $\log(n)$ factor, cannot be improved, in general, see Theorem 6.
- Second, we show that for specific sub-classes of trees (see Definition 1 below) `treeSeg` does overcome the $1/n$ rate and even yields a perfect segmentation with high probability, see Theorem 7.

The minimax optimal change-point estimation rate for an independent Bernoulli signal with piecewise constant success probabilities is of order $1/n$, see (1, 4). Thus, the first result in Theorem 6 shows that, for an arbitrary tree $T$, the additional tree information beyond the leaf ordering cannot improve the segmentation rate, in general. On the other hand, the second result in Theorem 7 shows that when the tree structure is sufficiently regular, the $1/n$ lower bound can, indeed, be improved. Hence, `treeSeg` efficiently explores the full information of the tree $T$ and outperforms regular change-point estimation procedures also in terms of segmentation rates in such cases the tree structure allows this. The overall improved segmentation performance of `treeSeg` compared to simple change-point procedures, such as (1), can also be observed empirically. We demonstrate this in a small simulation experiment in Figure S10.

In the following, let $\tau_1, \ldots, \tau_K \subset \{1, \ldots, n\}$ be the change-points of the true signal $p$, that is, $p_{\tau_i} \neq p_{\tau_i+1}$ and $\hat{\tau}_1, \ldots, \hat{\tau}_{\hat{K}} \subset \{1, \ldots, n\}$ the change-points of $\hat{p}$. Define the accuracy of the estimated change-point locations as

$$d(p, \hat{p}) := \max_{i=1,\ldots,K} \min_{j=1,\ldots,\hat{K}} \frac{1}{n} |\tau_i - \hat{\tau}_j| . \qquad [1]$$

Recall that Theorem 5 implies that for every change-point of $p$ there exists a change-point of $\hat{p}$ within a neighborhood of order at most $\log(n)$. More precisely, from Theorem 5 we obtain that for any fixed minimal scale $\lambda$, minimal success probability difference $\delta$, confidence level $\alpha$, and $p \in \mathcal{S}_{\lambda,\delta}$ it holds true that

$$d(p, \hat{p}) \leq C_3 \frac{\log(n)}{n} \qquad [2]$$

with probability at least $1 - C_1 e^{-C_2 n}$, where $C_1, C_2, C_3$ are positive constants that only depend on $\alpha, \lambda, \delta$. Also, recall that it follows from Theorem 2 that $\hat{k} = k$ with high probability whenever $\alpha$ is sufficiently small. The following theorem shows that this upper bound in [2] cannot be improved for a general tree $T$, possibly up the $\log(n)$ factor. To this end, let $\mathcal{S}_{\lambda,\delta}(k)$ denote the set of signals $p \in \mathcal{S}_{\lambda,\delta}$ with $k$ active nodes.

*Theorem 6.* For fixed minimal scale $\lambda \in (0, 1)$ and minimal success probability difference $\delta \in (0, 1)$, it holds true that there exists a tree $T$ with $k$ active nodes such that

$$\inf_{\substack{\hat{p} \\ \hat{k}=k}} \sup_{p \in \mathcal{S}_{\lambda,\delta}(k)} E_p\left(d(p, \hat{p})\right) \geq \frac{C}{n},$$

[*] Note that the change-points do not uniquely determine the active nodes, as an active node may induce one or two change-points depending on its location and thus, there can be a different number of active nodes that induce the same change points.

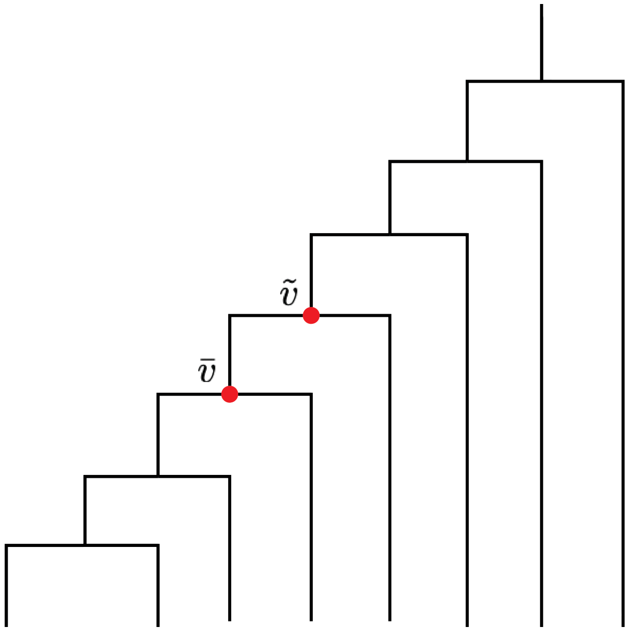


**Fig. S5.** Example of the tree $T^\star$ and nodes $\tilde{v}, \bar{v}$ for $n = 8$ as in the proof of Theorem 6.

where $C$ is a positive constant that only depends on $\delta$.
Theorem 6 shows that up to constants and possibly the log-factor, it is not possible to improve the $\log(n)/n$ rate as in [2] obtained from Theorem 5.

The proof of Theorem 6 follows directly by the observation that there always exist a tree (such as in Figure S5) which does not impose any additional restriction on the segmentation besides the overall leaf ordering. That means, the totally ordered change-point setting is just a special case within the class of all tree structures and hence, any lower bound for the change-point setting also applies for general tree structures. For sake of completeness, we give the explicit proof of Theorem 6 below.

*Proof of Theorem 6.* Let $T_n^\star$ denote a tree with $n$ leaf nodes as in Figure S5 and assume that $k = 1$, that is, there is a single active node $v$ which segments the leaf nodes into two segments. Note that the signal space $\mathcal{S}_{\lambda,\delta}(k)$ depends on the given tree $T$, so that we may write $\mathcal{S}_{\lambda,\delta}(k, T)$. Let $\bar{v}$ and $\tilde{v}$ be two inner nodes in $T_n^\star$, such that $\bar{v}$ is a direct offspring of $\tilde{v}$, that is, $\bar{v} \in \text{off}(\tilde{v})$, see Figure S5. Let $\bar{p}$ and $\tilde{p}$ be success probability signals of length $n$ with a single active node $\bar{v}$ and $\tilde{v}$, respectively, such that $\bar{p}|_{\text{Off}(\bar{v})} = \tilde{p}|_{\text{Off}(\tilde{v})} = p_0 + \delta$ and $\bar{p}|_{\text{Off}(\bar{v})^c} = \tilde{p}|_{\text{Off}(\tilde{v})^c} = p_0$, for some $p_0 \in (0, 1-\delta)$, where $\text{Off}(\bar{v})^c = \{1, \ldots, n\} \setminus \text{Off}(\bar{v})$, see Figure S5.

Clearly, for any arbitrary signal $p$ with a single active node $v$ (recall that we assume $k = 1$), it must hold true that the corresponding change-point of $p$ has distance at least 1 to either of the change-points of $\bar{p}$ or $\tilde{p}$, that is,

$$
\begin{aligned}
& n\, d(\bar{p}, p) + n\, d(\tilde{p}, p) = |\tau_1 - \bar{\tau}_1| + |\tau_1 - \tilde{\tau}_1| \geq 1 \\
\Leftrightarrow \quad & d(\bar{p}, p) + d(\tilde{p}, p) \geq 1/n.
\end{aligned} \qquad [3]
$$

Let $i^\star$ be the single leaf index in $T^\star$ which is an offspring of $\tilde{v}$ but not of $\bar{v}$, see Figure S5. Let $f_{\bar{p}}(Y)$ and $f_{\tilde{p}}(Y)$ denote the data likelihood under signal $\bar{p}$ and $\tilde{p}$, respectively. Then we have that

$$
\frac{f_{\tilde{p}}(Y)}{f_{\bar{p}}(Y)} = \frac{(p_0+\delta)^{Y_{i^\star}}(1-p_0-\delta)^{1-Y_{i^\star}}}{p_0^{Y_{i^\star}}(1-p_0)^{1-Y_{i^\star}}} =: Z,
$$

such that under model $\bar{p}$ we have that

$$
Z = \begin{cases} \frac{p_0+\delta}{p_0} & \text{with probability } p_0 \\ \frac{1-p_0-\delta}{1-p_0} & \text{with probability } 1-p_0. \end{cases}
$$

Note that

$$
E_{\bar{p}}\left(\min(1, Z)\right) = 1 - p_0 - \delta + p_0 = 1 - \delta.
$$

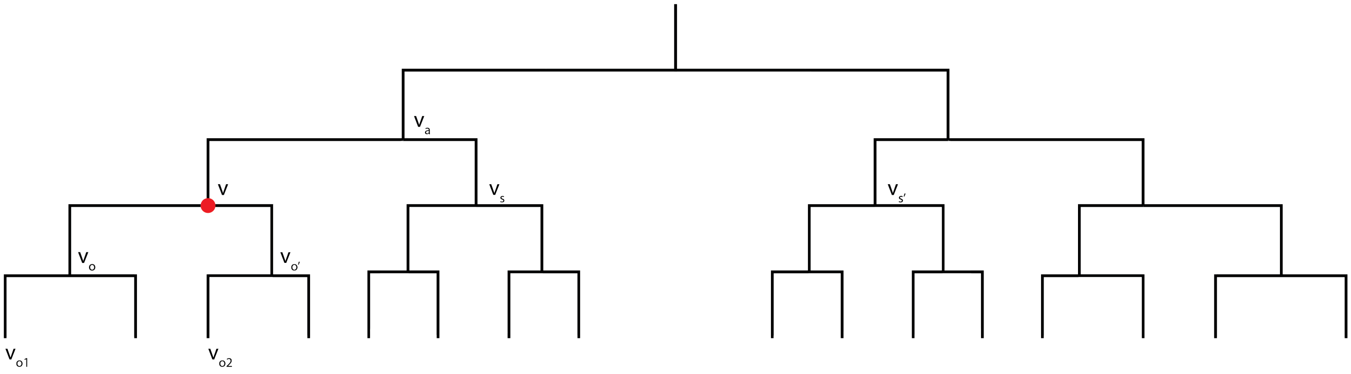


**Fig. S6.** A perfect binary tree with $n = 8$ leaf nodes. Illustrative example of notation $v, v_a, v_s, v_{s'}, v_o, v_{o'}, v_{o1}, v_{o2}$ as in the proof of Theorem 7.

Thus, it follows that

$$\begin{aligned}\sup_T \sup_k \inf_{\substack{\hat{p}\\ \hat{k}=k}} \sup_{p\in\mathcal{S}_{\lambda,\delta}(k)} E_p\left(d(p,\hat{p})\right) &\geq \inf_{\substack{\hat{p}\\ \hat{k}=1}} \sup_{p\in\mathcal{S}_{\lambda,\delta}(1,T^\star)} E_p\left(d(p,\hat{p})\right)\\ &\geq \frac{1}{2}\inf_{\hat{p}} E_{\bar{p}}\left(d(\bar{p},\hat{p})\right) + E_{\tilde{p}}\left(d(\tilde{p},\hat{p})\right)\\ &= \frac{1}{2}\inf_{\hat{p}} E_{\bar{p}}\left(d(\bar{p},\hat{p}) + d(\tilde{p},\hat{p})Z\right)\\ &\geq \frac{1}{2}\inf_{\hat{p}} E_{\bar{p}}\left(\left(d(\bar{p},\hat{p}) + d(\tilde{p},\hat{p})\right)\min(1,Z)\right)\\ &\geq \frac{1}{2n}\inf_{\hat{p}} E_{\bar{p}}\left(\min(1,Z)\right)\\ &= \frac{1-\delta}{2n}\end{aligned}$$

□

In the following we complement Theorem 6 with a result of a more positive flavor. Although, the tree structure cannot improve the segmentation rate in general, when one imposes additional assumptions on the tree, `treeSeg` is able to overcome the $1/n$ lower bound and even obtains a perfect segmentation with high probability.

*Definition 1.* Let $\gamma, \lambda \in (0,1)$, $k \in \mathbb{N}$ . We denote a binary tree $T$ with $n$ leaf nodes and root $v_0 = 2n-1$ as *$\gamma$-spreading* at minimal scale $\lambda$, if for any collection of inner nodes $v_1, \ldots, v_k \in V_I$ that induce a segmentation with minimal scale $\lambda$, there exist disjoint intervals $\mathcal{I}_1^1, \mathcal{I}_1^2, \ldots, \mathcal{I}_m^1, \mathcal{I}_m^2 \subset \{1,\ldots,n\}$ of minimal length $\gamma\lambda$ such that

1. for all $i = 1,\ldots,m$, $l = 1,2$ we have that $\mathcal{I}_i^l$ belongs to the influence region of exactly one node $v_j$, that is, $\mathcal{I}_i^l \subset \mathrm{IR}(v_j)$, for some $j = 0,\ldots,k$,
2. for all $i = 1,\ldots,m$ we have that $\mathcal{I}_i^1$ and $\mathcal{I}_i^2$ belong to influence regions of different nodes $v_{j_1}, v_{j_2}, j_1, j_2 = 0,\ldots,k$, that is, $\mathcal{I}_i^1 \subset \mathrm{IR}(v_{j_1})$ and $\mathcal{I}_i^2 \subset \mathrm{IR}(v_{j_2})$ with $v_{j_1} \neq v_{j_2}$,

and for any other inner nodes $\{\tilde{v}_1,\ldots,\tilde{v}_k\}$ that induce a different segmentation there exists an interval index $i = 1,\ldots,m$ and a node $j = 0,\ldots,k$ such that $\mathcal{I}_i^1$ and $\mathcal{I}_i^2$ are both in the influence region of $\tilde{v}_j$, that is $\mathcal{I}_i^1, \mathcal{I}_i^2 \subset \mathrm{IR}(\tilde{v}_j)$, with root node $\tilde{v}_0 = 2n-1 = v_0$.

Definition 1 captures precisely what is needed for `treeSeg` to obtain perfect segmentation with high probability uniformly among all possible active nodes in the tree $T$, given a minimal scale $\lambda$. The following example shows that $\gamma$-spreading trees can be seen as a generalization of perfect trees, see Figure S6.

*Example 1. (Perfect trees)* A simple example of a $\gamma$-spreading tree is a binary perfect tree, that is, a tree in which all internal nodes have two children and all leaf nodes are at the same level. Consider the case $k = 1$. Let $v = v_1 \in V_I$ be some inner node of $T$ which induces a segmentation of minimal scale $\lambda$. Assume that $v$ is at level $L$, that is, $|\mathrm{IR}(v)| = |\mathrm{Off}(v)| = n/2^L \geq n\lambda$. See Figure S6 for an example. Assume that $L \geq 2$ (the case where $L = 1$ is similar). Let $v_o, v_{o'}$ be the two direct offspring nodes of $v$ at level $L+1$, that is $\{v_o, v_{o'}\} = \mathrm{off}(v)$. Let $v_{o1} \in \mathrm{off}(v_o)$ and $v_{o2} \in \mathrm{off}(v_{o'})$ be two offspring nodes at level $L+2$ of $v_o$ and $v_{o'}$, respectively, see Figure S6. Let $v_a$ be the direct ancestor node of $v$ and let $v_s$ be the (unique) sibling node of $v$, that is, $\{v, v_s\} = \mathrm{off}(v_a)$. Let $v_{s'} \notin \{v, v_s\}$ be any other inner node at level $L$. Note that, because the tree $T$ is a perfect binary tree, we have that

$$|\mathrm{Off}(v_{o1})| = |\mathrm{Off}(v_{o2})| = |\mathrm{Off}(v)|/4 \geq n\lambda/4$$

and

$$|\mathrm{Off}(s)| = |\mathrm{Off}(s')| = |\mathrm{Off}(v)| \geq n\lambda.$$

Define

$$\mathcal{I}_1^1 = \mathrm{Off}(v_{o1}),\ \mathcal{I}_1^2 = \mathrm{Off}(v_s)$$
$$\mathcal{I}_2^1 = \mathrm{Off}(v_{o2}),\ \mathcal{I}_2^2 = \mathrm{Off}(v_{s'})$$

See Figure S6 for an example. Clearly, the intervals $\mathcal{I}_i^l$ fulfill conditions 1. and 2. of Definition 1. Let $\tilde{v} \neq v \in V_I$ be any other inner node of $T_n$. We distinguish three different cases.

- If $\tilde{v}$ *is not an offspring of* $v_a$, then $\mathcal{I}_1^1, \mathcal{I}_1^2$ are both either in the influence region of $\tilde{v}$, that is, $\mathcal{I}_1^1, \mathcal{I}_1^2 \subset \mathrm{IR}(\tilde{v})$, or of the root $\tilde{v}_0 = 2n-1$, that is, $\mathcal{I}_1^1, \mathcal{I}_1^2 \subset \mathrm{IR}(\tilde{v}_0)$.
- If $\tilde{v}$ *is an offspring of* $v$, say $\tilde{v}$ is equal to or an offspring of $v_o$ (or $v_{o'}$, respectively), then $\mathcal{I}_2^1, \mathcal{I}_2^2$ (or $\mathcal{I}_1^1, \mathcal{I}_1^2$, respectively) are both in the influence region of the root $v_0$, that is, $\mathcal{I}_2^1, \mathcal{I}_2^2 \subset \mathrm{IR}(\tilde{v}_0)$ (or $\mathcal{I}_1^1, \mathcal{I}_1^2 \subset \mathrm{IR}(\tilde{v}_0)$, respectively).
- If $\tilde{v}$ *is either equal to or an offspring of* $v_s$, then $\mathcal{I}_2^1, \mathcal{I}_2^2$ are both in the influence region of the root $\tilde{v}_0 = 2n-1$, that is, $\mathcal{I}_2^1, \mathcal{I}_2^2 \subset \mathrm{IR}(\tilde{v}_0)$.

This shows that a perfect tree $T$ is 0.25-spreading for a single active node $k = 1$.

*Theorem 7.* Fix a minimal scale $\lambda > 0$, minimal success probability difference $\delta > 0$, confidence level $1-\alpha \in (0,1)$, and spreading parameter $\gamma > 0$. Then uniformly over all $\gamma$-spreading trees $T$ and signals $p \in \mathcal{S}_{\lambda,\delta} = \mathcal{S}_{\lambda,\delta}(T)$, for the `treeSeg` estimate $\hat{p}$, conditioned on $\hat{k} = k$, it holds true that

$$d(p, \hat{p}) = 0,$$

with probability at least $1 - C_1\, e^{-C_2\, n}$, where $C_1, C_2$ are positive constants which only depend on $\alpha, \lambda, \delta, \gamma$.
The proof of Theorem 7 follows directly from the definition of $\gamma$-spreading (see Definition 1) together with power considerations of the LR test, analog as discussed in the proof of Theorem 2, see Section C.

Theorem 7 shows that when the tree $T$ is sufficiently regular (as in the definition of $\gamma$-spreading of which perfect trees are a special case), then with high probability `treeSeg` estimates the locations of the active nodes exactly, overcoming the $1/n$ lower bound from the simple change-point setting (without tree structure).

Moreover, `treeSeg`'s confidence sets $\mathcal{C}_{1-\alpha}$ for the active nodes directly capture this exact recovery property. That means, when a tree satisfies the conditions of Theorem 7 then with high probability $|\mathcal{C}_{1-\alpha}| = \hat{k} = k$, that is, $\mathcal{C}_{1-\alpha}$ will only contain the true active nodes $\mathcal{C}_{1-\alpha} = \{v_1, \ldots, v_k\}$ so that exact recovery is directly obtainable from `treeSeg`'s output.

**E. Confidence statements with random branch swapping.** Recall that the `treeSeg` algorithm does, in general, depend on the specific ordering of the leaves induced by the tree structure. This ordering, however, is not uniquely determined by the tree itself. For every inner node of the tree one can swap the two offspring nodes to generate a new ordering of the leaves which represents the tree equally well.

Our theoretical guarantees hold true for any (arbitrary) such ordering, as long as this ordering was chosen a-priori, independent of the data $Y$. In a simulation study in Section B we also show that `treeSeg`'s overall segmentation among random swapping of tree branches remains reasonably stable.

Clearly, if one chooses the specific branch ordering based on the observations $Y_i$, theoretical guarantees may be violated, as this essentially amount to *p-value hunting*. This problem is not intrinsic to `treeSeg`, but is a universal problem of statistics whenever there are several tests for the same hypothesis problem available. Deciding on one particular branch ordering amounts to deciding on one of the particular tests, which opens the door for so called *p-value lottery/ p-value hunting*.

A natural approach to avoid this arbitrary choice is to apply `treeSeg` repetitively to $B$ different leaf-orders, obtained via randomly swapping the branches of the tree, and then to aggregate the results. When $B$ is sufficiently large, this will make the analysis (at least approximately) independent of any specific branch swapping.

In the following we describe a simple-to-implement way for such an aggregation which preserves `treeSeg`'s overall confidence statements, by adapting an approach from (5) to a similar problem for p-values in high-dimensional linear regression.

Fix some confidence level $\alpha \in (0,1)$. For $b = 1, \ldots, B$ randomly obtained branch swapped version of the tree $T$ and data $Y$, which correspond to different orderings of $Y_i$. Apply `treeSeg` at level $\alpha/2$, to obtain estimates for the number of active nodes $\hat{k}^b(q_{1-\alpha/2})$, confidence sets for the active nodes $\mathcal{C}^b_{1-\alpha/2}$, and confidence bands for the success probabilities $\underline{p}^b_{1-\alpha/2}, \overline{p}^b_{1-\alpha/2}$, for each $b = 1, \ldots, B$. Aggregate these results as follows:

$$\begin{aligned} \hat{k} &:= \mathrm{median}_{b=1,\ldots,B}\hat{k}^b(q_{1-\alpha/2}), \\ \mathcal{C} &:= \{i \in V \ :\ \frac{1}{B}\sum_{b=1}^{B} \mathbb{1}_{i \in \mathcal{C}^b_{1-\alpha/2}} \geq 0.5\}, \\ (\underline{p})_i &:= \mathbf{median}_{b=1,\ldots,B}(\underline{p}^b_{1-\alpha/2})_i, \qquad \text{for } i = 1,\ldots,n \\ (\overline{p})_i &:= \mathbf{median}_{b=1,\ldots,B}(\overline{p}^b_{1-\alpha/2})_i, \qquad \text{for } i = 1,\ldots,n \end{aligned} \qquad [4]$$

The choice of the median together with the level correction $\alpha/2$ in [4] is somewhat arbitrary and can be replaced with any other quantile $\gamma \in (0,1)$ together with level correction $\gamma\alpha$, see (5) for details.

It is easy to check that for the aggregated treeSeg results in [4] all confidence statements hold true at level $\alpha$. We illustrate this for the estimated number of active nodes $\hat{k}$ and for the confidence set $\mathcal{C}$, the derivation for the confidence band $\overline{p}, \underline{p}$ is analog. We have that

$$\mathbf{P}\left(\hat{k} > k\right) = \mathbf{P}\left(\text{median}_{b=1,\dots,B}\ \hat{k}^b(q_{1-\alpha/2}) > k\right) = \mathbf{P}\left(\frac{1}{B}\sum_{b=1}^{B} \mathbb{1}_{\hat{k}^b(q_{1-\alpha/2})>k} > 0.5\right) \leq 2\ E\left(\frac{1}{B}\sum_{b=1}^{B} \mathbb{1}_{\hat{k}^b(q_{1-\alpha/2})>k}\right) \leq \alpha,$$

where the first inequality follows from Markov's inequality and the second inequality follows because for all individual random branch swaps $b$ treeSeg guarantees that $\mathbf{P}\left(\hat{k}^b(q_{1-\alpha}) > k\right) \leq \alpha$. Similar, one obtains that

$$\begin{aligned}
\mathbf{P}\left(\exists_{i=1,\dots,k}\ v_i \notin \mathcal{C}\right) &= \mathbf{P}\left(\max_{i=1,\dots,k} \frac{1}{B}\sum_{b=1}^{B} \mathbb{1}_{v_i \notin \mathcal{C}^b_{1-\alpha/2}} \geq 0.5\right) \\
&\leq \mathbf{P}\left(\frac{1}{B}\sum_{b=1}^{B} \max_{i=1,\dots,k} \mathbb{1}_{v_i \notin \mathcal{C}^b_{1-\alpha/2}} \geq 0.5\right) \\
&\leq 2E\left(\frac{1}{B}\sum_{b=1}^{B} \max_{i=1,\dots,k} \mathbb{1}_{v_i \notin \mathcal{C}^b_{1-\alpha/2}}\right) \\
&= 2E\left(\frac{1}{B}\sum_{b=1}^{B} \mathbb{1}_{\exists_{i=1,\dots,k}\ v_i \notin \mathcal{C}^b_{1-\alpha/2}}\right) \leq \alpha,
\end{aligned}$$

where the first inequality follows because the maximum over sums is smaller or equal to the sums over maxima, the second inequality follows from Markov's inequality, and the last inequality from the coverage probability of each $\mathcal{C}^b_{1-\alpha/2}$ individually, that is $\mathbf{P}\left(\exists_{i=1,\dots,k}\ v_i \notin \mathcal{C}^b_{1-\alpha}\right) \leq \alpha$.

**F. Noisy tree structure.** The treeSeg algorithm takes as input a given tree structure $T$ with $n$ leaf nodes as well as the observations of interest $Y_1, \dots, Y_n$. Thereby, the tree $T$ defines some neighborhood structure between the observations $Y_1, \dots, Y_n$. The question which treeSeg answers, is whether the distribution of observation $Y_i$ depends on its relative location within the neighborhood structure defined by $T$.

Clearly, in many (if not most) applications, the tree $T$ is itself just a noisy surrogate of the actual neighborhood structure of interest. For example, a phylogenetic tree obtained from genetic sequencing data, will usually not exactly correspond to the true ancestral chart of the individuals under consideration.

Similar, for one and the same observations $Y_1, \dots, Y_n$ there might be different trees that correspond to different neighborhood structures of interest. Consider, for example, the three plots in Figure S7. They all show the same observations $Y_1, \dots, Y_{180}$, simulated from independent Bernoulli observations with success probability as shown as the black dashed line. However, the trees which define their neighborhood structures are different. Arguably, in the very right plot there is no obvious dependence between the tree structure and the distribution of the observations $Y_i$. In the other two plots, however, the neighborhood structure is significantly different and the success probability strongly depends on particular sub-branches in the tree.

The example in Figure S7 was generated as follows. We considered $n = 180$ Bernoulli random variables $Y_1, \dots, Y_{180}$ such that

$$\begin{aligned}
Y_1, \dots, Y_{60} &\overset{\text{i.i.d.}}{\sim} B(0.1) \\
Y_{61}, \dots, Y_{120} &\overset{\text{i.i.d.}}{\sim} B(0.5) \\
Y_{121}, \dots, Y_{180} &\overset{\text{i.i.d.}}{\sim} B(0.9).
\end{aligned} \qquad [5]$$

Then, we simulated $n = 180$ independent multivariate Gaussian random variables such that

$$\begin{aligned}
X_1, \dots, X_{60} &\overset{\text{i.i.d.}}{\sim} N\left((0,0)^\top, \sigma\ \text{Id}_2\right) \\
X_{61}, \dots, X_{120} &\overset{\text{i.i.d.}}{\sim} N\left((1,0)^\top, \sigma\ \text{Id}_2\right) \\
X_{121}, \dots, X_{180} &\overset{\text{i.i.d.}}{\sim} N\left((0,1)^\top, \sigma\ \text{Id}_2\right).
\end{aligned} \qquad [6]$$

For $\sigma = 0.01, 0.07$ and 1, we applied hierarchical clustering to the observations $X_i$ to obtain a tree structure as shown in Figure S7.

For $\sigma = 0.01$ the tree $T$ perfectly aligns observations from the same group into a joint sub-branch of the tree. Hence, treeSeg perfectly recovers this segmentation which is represented by the tree. On the other extreme, for $\sigma = 1$, the tree essentially does not capture any neighborhood structure from the original grouping of the observations. Hence, treeSeg, which analyses the data conditioned on this particular tree $T$, does not estimate any active nodes, hence, does not report any dependence between the tree and the observational distribution. Given that this tree essentially does not contain any dependence structure, this is a very reasonable results.

In summary, we stress that `treeSeg` does not make any assumptions on how the particular tree $T$ was generated. It takes the tree $T$ as given and performs its analysis conditioned on $T$. Only this enables `treeSeg`'s wide applicability to various settings, where trees might be generated in very different ways.

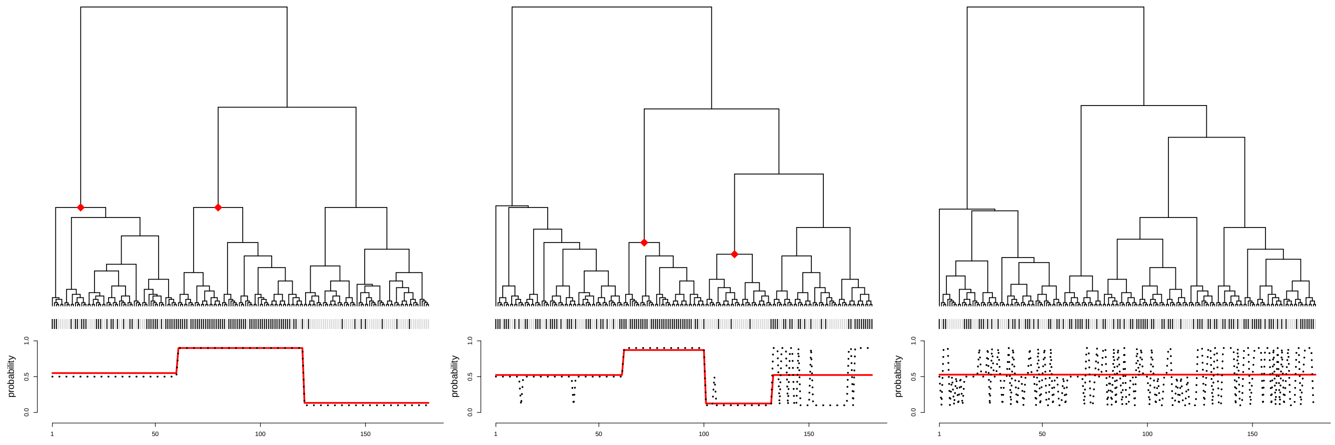

**(a)** $\sigma = 0.01$ **(b)** $\sigma = 0.07$ **(c)** $\sigma = 1$

**Fig. S7.** Trees obtained via clustering of $X_i$'s as in [6] together with Bernoulli observations at the leaves of the tree as in [5]. The success probability from which $Y_i$ was simulated is shows as black dashed line. The estimate $\hat{p}$ obtained from `treeSeg` with $\alpha = 0.1$ is shown in red together with its estimated active nodes in the tree.

### 2. Simulation results

In the following we report on simulation results for the `treeSeg` algorithm. We have performed two sets of simulation studies. In the first study we assessed the power of the method in detecting clades on the tree with distinct phenotype distributions. In the second study we assessed the impact of the tree branch ordering on the outcome of the method.

#### A. Power of the method in detecting clades with distinct phenotype distributions.

***A.1. Single active node is simulated:.*** First we want to study the detection power of `treeSeg` of active components on the tree, that is, subtrees where the underlying success probability of the binary response variables at the leaves differs from the rest of the tree. The power is dependent on two parameters, the number of samples contained in the subtree, the larger the subtree, the easier it is to be detected. Similarly, the larger the difference between success probabilities in active sub-tree and rest of the tree is, the easier it is to detect this sub-tree. Therefore, we can quantify the difficulty level of the detection problem via two parameters: First, the size of the smallest subgroup, which we denote by $\lambda$. Second, the size of the smallest pairwise difference of success probabilities, which we denote by $\delta$. To study the ability of `treeSeg` to detect an active subgroup we consider the following setup:

We fix the sample size $n = 200$, fix a confidence level $\alpha = 0.1$. Then, for each $\delta = 0.1, 0.2, \ldots, 1$ and each $\lambda = 5, 15, 25, 35, \ldots, 95$ we do the following:

1. We generate a random binary tree with $n$ leaves.
2. We select a random inner node which has exactly $\lambda$ offspring nodes (if this does not exist we go back to 1.).
3. We select randomly two probabilities $p_1, p_2$ with $|p_1 - p_2| = \delta$.
4. We draw binary observations with success probability $p_1$ for the offspring of the selected node and with success probability $p_2$ for other nodes.
5. We apply the `treeSeg` algorithm.

Then, we repeat until convergence of the reported summary statistics (we found $1,000$ repetitions to be sufficient).

Figure S11 shows the average number of detected active nodes for different values of $\delta$ and $\lambda$. Note that, in this setting, the true number of active nodes is one. One can see that for reasonable values of $\lambda$ and $\delta$ `treeseg` detects an active sub-group with very high probability and thus, reveals a strong detection power. In particular, Figure S11 confirms that `treeseg` reliably controls the probability of over estimation. Indeed, we found that for all combinations of $\delta$ and $\lambda$ the empirical probability of overestimation was even smaller than the nominal level $\alpha = 0.1$. This reveals that active components, which get detected by `treeseg`, are indeed present in the signal with very high probability. More importantly, this probability guarantee can be made explicit via the level $\alpha$ (in this setting $\alpha = 0.1$).

Figure S12 gives an insight on the accuracy of the estimated active node (in an detection event). It displays the average distance of the estimated active node in the tree to the true active node. Thereby, as a distance function we considered the

 Merle Behr, M. Azim Ansari, Axel Munk, and Chris Holmes

minimum number of edges to be traversed to reach one node from other. One can see that for reasonable $\lambda$ and $\delta$ the estimated active node coincides with the true active node, which shows a high estimation accuracy of `treeseg`.

Finally, we report on the estimation accuracy for the leaf-success probabilities. Figure S13 shows the mean of absolute error (MAE) of of the estimated success probabilities for different $\lambda$ and $\delta$. Note that MAE are small in general, revealing a high accuracy of `treeseg` for success probability estimation. For small $\delta$, MAE slightly increases as $\lambda$ increases. This is because when $\delta$ is small, the active node cannot be detected reliably and the smaller $\lambda$, the better one can approximate the true success probabilities via a constant. When $\delta$ is reasonably large, `treeseg` reliably detects the active node and MAE naturally increases as the $\lambda$ increases.

***A.2. No active node is simulated:.*** First, we consider the situation where the true generating model has no active component, that is all responses at the tree leaves have the same success probability. The following setup is considered. We fix the sample size $n = 200$ and a confidence level $\alpha = 0.1$.

1. We generate a random binary tree with $n$ leaves.
2. We select randomly a single success probability $p_1$.
3. We draw $n$ binary observations with success probability $p_1$.
4. We apply the `treeseg` algorithm.

Then, we repeat until convergence of the reported summary statistics (we found $1,000$ repetitions to be sufficient). We found that in 98.6% of cases `treeseg` agreed with simulation truth. Note that this is even higher than our theoretical guarantee of $1 - \alpha = 0.9$. This again shows that if `treeseg` does detect an active component, this is very strong evidence that this is indeed present in the signal.

***A.3. Two active nodes are simulated.*** In this study, we considered generating models with two active nodes. More precisely, we considered the following setup. We fix the sample size $n = 200$, fix a confidence level $\alpha = 0.1$. Then, for each $\delta = 0.1, 0.2, \ldots, 0.5$ and each $\lambda = 5, 15, 25, 35, \ldots, 65$ we do the following:

1. We generate a random binary tree with $n$ leaves.
2. We select two random inner node with each having exactly $\lambda$ offspring nodes (if this does not exist we go back to 1.).
3. We select randomly three probabilities $p_1 < p_2 < p_3$ with $|p_1 - p_2| = |p_2 - p_3| = \delta$.
4. We draw binary observations with success probability $p_i$ the respective sub-groups (assigning randomly $p_1, p_2, p_3$ to the sub-groups).
5. We apply the `treeseg` algorithm.

Then, we repeat until convergence of the reported summary statistics (we found $1,000$ repetitions to be sufficient).

Figure S14 shows the average number of detected active nodes for different values of $\delta$ and $\lambda$. Note that, in this setting, the true number of active nodes is two. One can see that for reasonable values of $\lambda$ and $\delta$ `treeseg` detects an active sub-group with high probability and thus, reveals a strong detection power. Compared to the setting with just a single active component, $\delta$ and $\lambda$ only need to be slightly larger to obtain detection of both active components simultaneously with high probability.

In particular, Figure S14 confirms that `treeseg` reliably controls the probability of overestimation. Again, for all combinations of $\delta$ and $\lambda$ the empirical probability of overestimation was even smaller than the nominal level $\alpha = 0.1$. This reveals that also when several active components are detected with very high probability they are indeed present in the data and this probability guarantee can be made explicit via the level $\alpha$ (in this setting $\alpha = 0.1$).

Figure S15 gives an insight on the accuracy of the estimated active nodes (in an detection event). It displays the average distance of the estimated active node in the tree which was closest to first true active node (for symmetry reasons the results for the second active nodes are the same). One can see that for reasonable $\lambda$ and $\delta$ the estimated active nodes are very close to the true active nodes, which shows a high estimation accuracy of `treeseg`.

For completeness, we also present simulations results for the MAE (analog to that in Figure S13) in Figure S16. They show a similar pattern as before with a decent estimation accuracy, in general.

**B. Impact of tree branching order on the estimation procedure.** Finally, we examine the impact of the tree branching order via simulations. Recall that the theoretical guarantees of `treeSeg` hold true for any specific leaf ordering, as long as this was selected independently of the data, that is, not as in a *p-value lottery.* In general, however, `treeSeg`'s output may differ for different orderings of the leaf nodes induced by the same tree. Recall the discussion in Section E.

For practical purposes, our current implementation of `treeSeg`[†] performs a standardized pre-ordering of the branches via the `reorder.phylo(..., order = "cladewise")` function of the `ape` R-package. In that way, results are independent of any specific ordering provided by the user. [‡]

[†] https://github.com/merlebehr/treeSeg

[‡] We want to note here, that this choice of cladewise ordering is rather arbitrary and only serves the purpose of standardization. In general, the *optimal* branch ordering (in terms of detection power) will depend on the signal, as well as on the data. Empirically we could not observe any *a-priori* best or worst ordering, independent of signal and data.

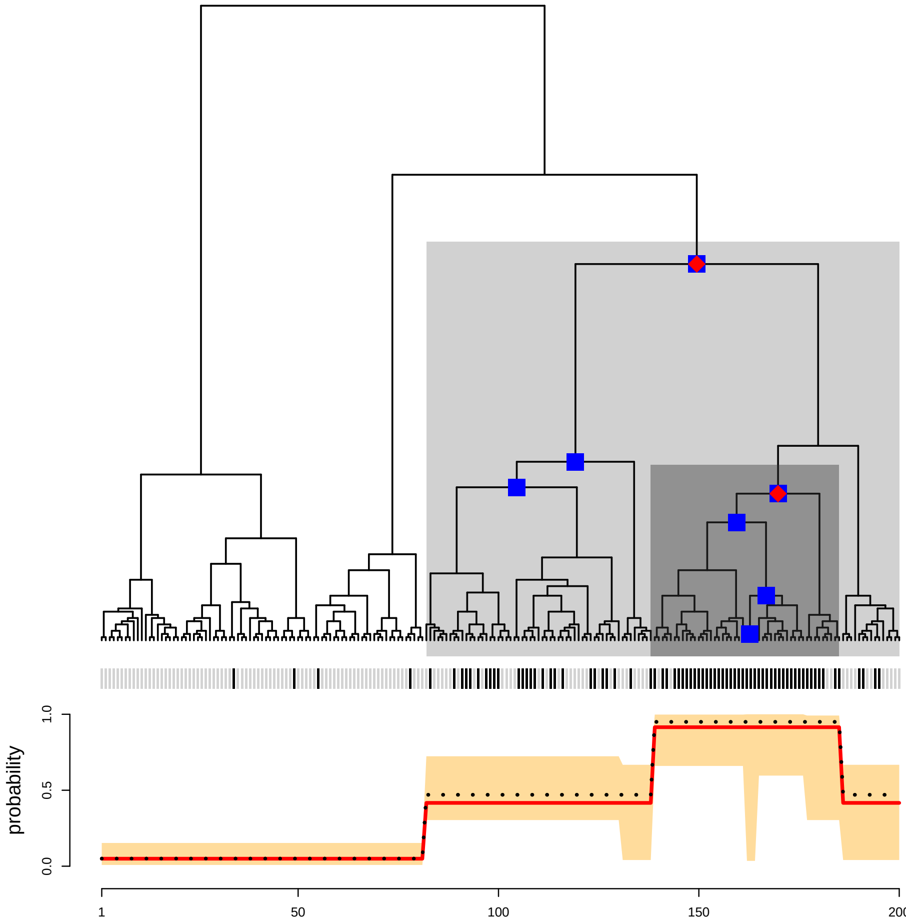


**Fig. S8.** The same setting as in Figure 1 but with the branches corresponding to the second active node and its sibling node swapped.

In order to still examine `treeSeg`'s behavior under different leaf orders, we also provide the option `tipOrder = unchanged`, which keeps the leaf order unchanged. As an example, reconsider the simulated observations as in Figure 1. The second active node induces a segment with 0.95 success probability at the very right of the leaf ordering. If one swaps the two branches that correspond to the direct ancestor of this active node, the same segment will appear more in the center, although the tree structure remains unchanged.

This is shown in Figure S8, together with `treeSeg`'s output. As can be seen, this branch swapping does not influence `treeSeg`'s estimated segmentation in this example. Only its confidence set $\mathcal{C}_{0.9}$ changes slightly. This is because for a different ordering, the tests that `treeSeg` applies, will differ slightly and thus, in principle may lead to slightly different results.

In the following, we show in a simulation study, that this stable behavior which we observe for the example in Figure 1 and S8, is the usual situation. As long as branch swapping is done randomly, `treeSeg`'s results typically do not change much.

Thus, in order to avoid *p-value hunting* we found it to be best practice to just fix some pre-specified ordering, e.g., cladewise as in our current implementation. Another approach would be via aggregating results over several random branch swaps, as discussed in Section E. However, the overall branch swapping stability of `treeSeg` observed in practice, indicates that the additional computational burden of the second approach might not always be justified.

To analyze the branch swapping effect, we will consider the following simulation setup. We fix the sample size $n = 200$ and the confidence level $\alpha = 0.1$. Further, in order to guarantee some detection, we fix a minimal $\lambda = 20$ and $\delta = 0.3$ (we later also consider other difficulty regimes via $\lambda, \delta$). Then we do the following:

1. We generate a random binary tree with $n$ leaves.
2. We select a random inner node with at least $\lambda$ offspring nodes (if not exists go back to 1.).
3. We select randomly two probabilities $p_1, p_2$ with $|p_1 - p_2| = \delta$.
4. Draw binary observations with success probability $p_1$ for the offspring of the selected node and with success probability $p_2$ for other nodes.
5. Then, for $N = 100$ repetitions

(a) With probability 1/2 independently swap the offspring of each branch of the tree.

(b) Apply `treeseg` algorithm to (swapped) data.

Figure S17 shows the histogram of empirical standard deviation among 100 randomly swapped leaf orderings for $1,000$ trees (generated as explained above). One can see that for most of the trees the standard deviation was zero, meaning that in all 100 branch swaps the number of detected active nodes was the same. Similar, Figure S18 shows the respective empirical 90% inter quantile distance, which shows for the majority of the $1,000$ trees at least 90 of the 100 randomly swapped leaf orderings resulted in the same number of estimated active nodes.

Also, in terms of estimation accuracy, branch swapping has very little influence. This can be seen in Figure S19, which shows that for the vast majority of trees the empirical standard deviation of distance between true and estimated active nodes among the 100 randomly swapped leaf orderings was almost zero. This shows that `treeseg` is very robust under possible branch swapping.

This robustness even becomes stronger in both, the very high and very low signal to noise ratio regimes. We repeated the experiment, but with minimal $\lambda = 50$ and minimal $\delta = 0.4$. Figure S20, S22, and S21 show the empirical standard deviation of number of active nodes, 90% inter quantile distance, and empirical standard deviation of distance between true and estimated active nodes. In this regime, for almost all of the $1,000$ trees branch swapping never influenced the algorithms outcome. The same holds true for the low signal to noise ratio regime, as seen in Figure S23, S25, and S24, where we considered a maximal $\lambda = 20$ and maximal $\delta = 0.2$. In summary, the simulations suggest that `treeseg` is quite robust under branch swapping events, in particular in low and high signal to noise regimes.

### 3. Application example on host specific epidemics of Salmonella bacteria

We also applied `treeSeg` to a data set on Salmonella bacteria from (6). The study considered 360 strains of Salmonella Typhimurium DT104 bacteria of which 113 were obtained from animal and 247 from human hosts. Typhimurium DT104 can infect both animals and humans, but it is unclear whether there are sublineages within DT104 that infect one host type more than the other and to what extent the epidemics in animals and humans are associated. A maximum likelihood phylogenetic tree was obtained from whole-genome sequencing data of the 360 strains, see (7) for details. Figure S9 shows `treeSeg`'s output for the phlyogenetic tree where the response variable is the host type (human vs. animal) from which the isolate was sampled. `treeSeg` finds two active nodes which divide the tree into three segments (two of the segments are non-contiguous given the tree leaf ordereing), that there were sub-lineages in the bacterial population with an increased restriction to infecting animals.

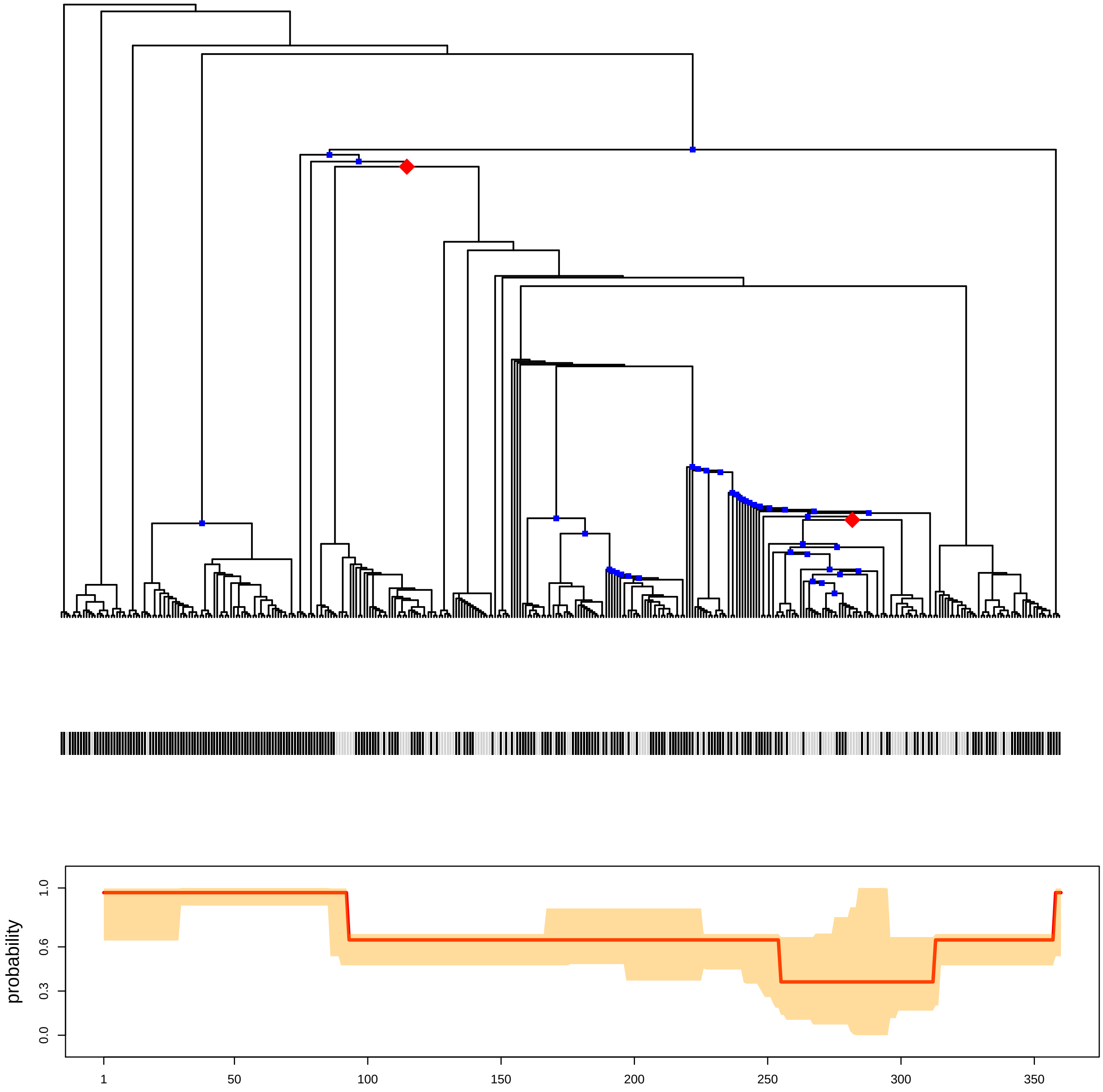


**Fig. S9.** Data example from (6) where a total of $360$ Salmonella enterica bacterial strains were studied. The tree shows a maximum-likelihood phylogeny based on whole genome sequencing of the strains. The color code at the leaves of the tree indicates whether the sample was isolated from human (black) or animal (gray) host. Using $\alpha = 0.1$, `treeSeg` estimates two active nodes. Localization of the first active node obtains a higher certainly compared to the second one, as seen in the confidence sets and bands.

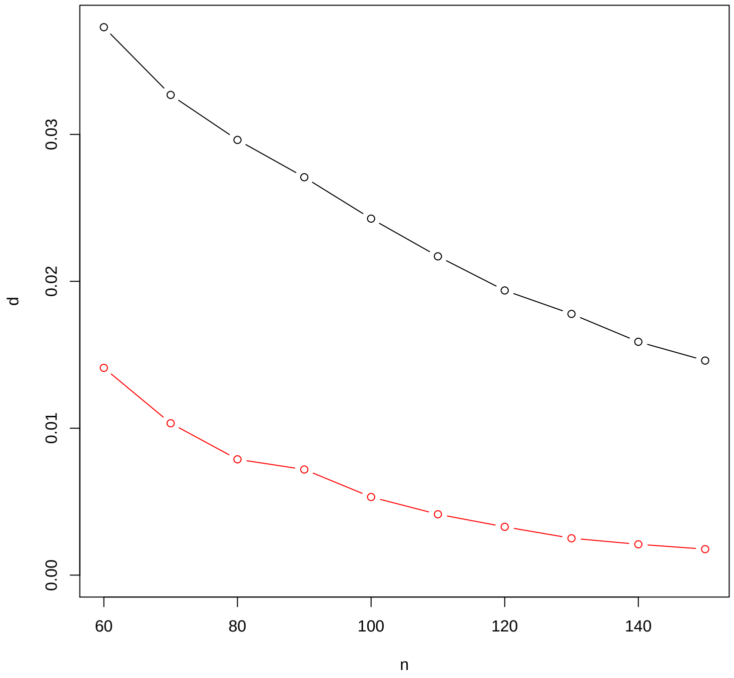


**Fig. S10.** Simulation example comparing change-point estimation accuracy of `treeSeg` and `SMUCE` (1). For $n = 60, 70, \dots, 150$ we simulated a random tree with a single active node splitting the leaf nodes into two parts of (approximately) equal size. For the two segments we generated Bernoulli random variables with success probability $0.1$ and $0.6$, respectively. The y-axis shows $d(p, \hat{p})$ as in [1] for `treeSeg` (red) and `SMUCE` (black), both with $\alpha = 0.1$, averaged over $10,000$ Monte-Carlo repetitions.

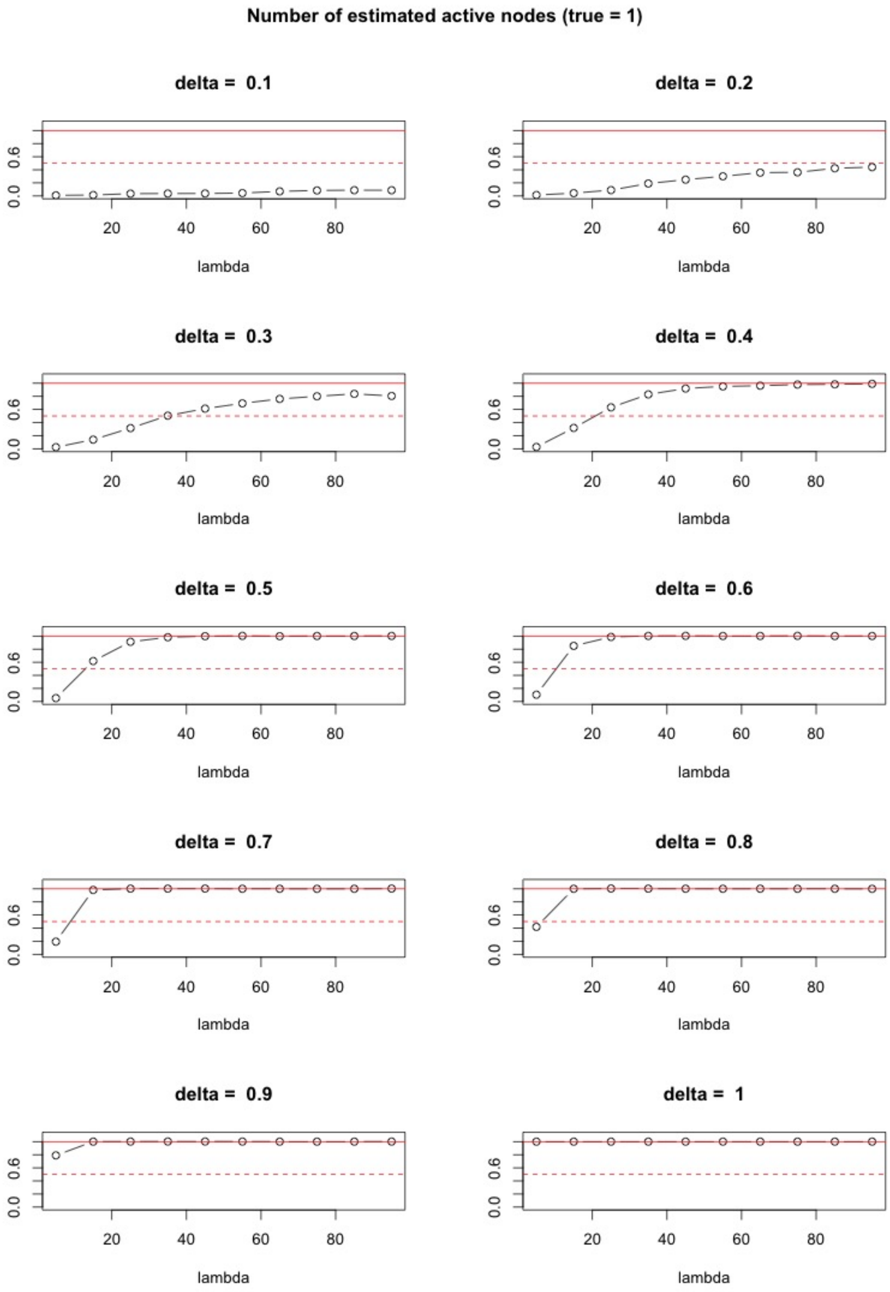


**Fig. S11.** Average number of detected active nodes for different minimal jump sizes $\delta$ and size of active sub-group $\lambda$ in simulations of Setting 1, with true number of active nodes equal to one.

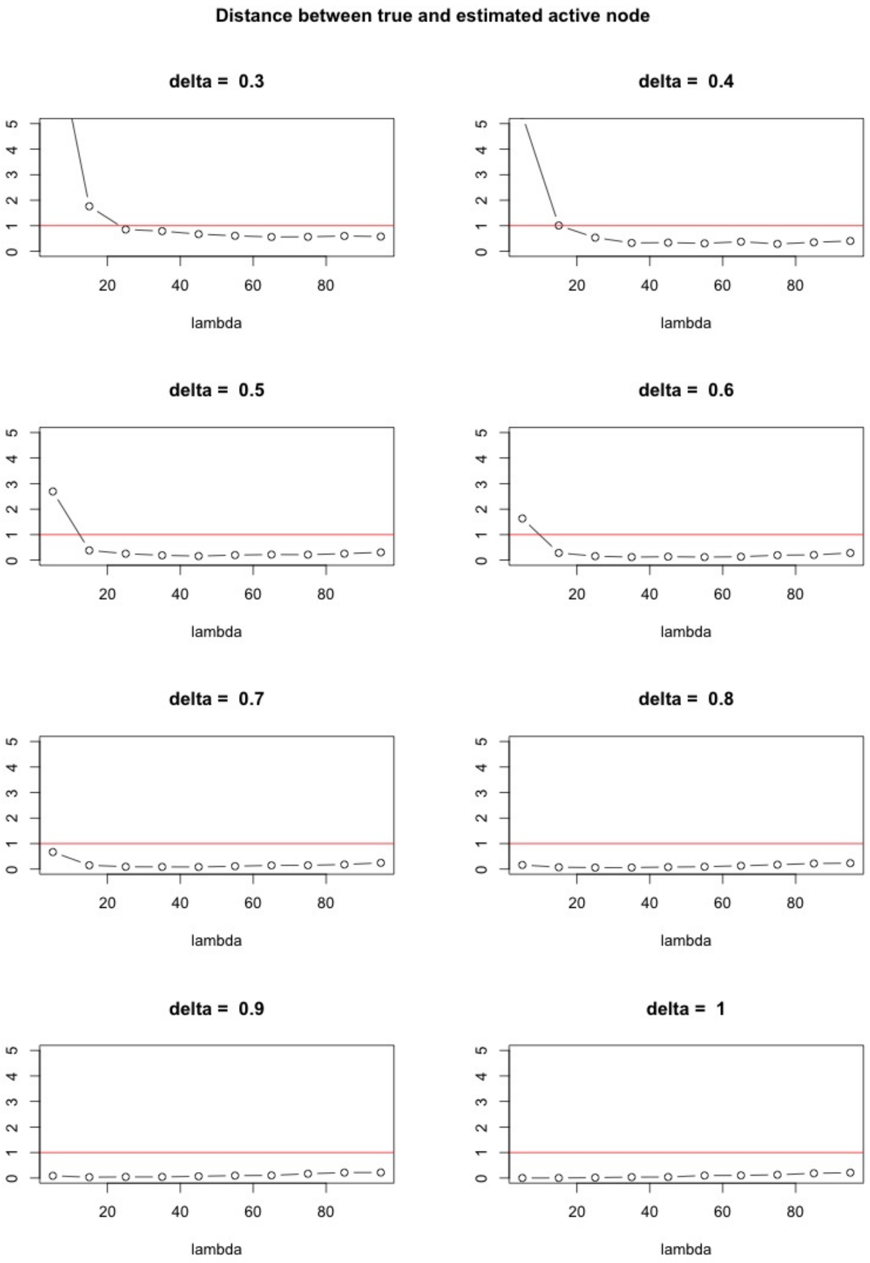


**Fig. S12.** Average distance of detected active node to the true active node (conditioned on detection event), with distance function being the minimum number of edges to be traversed to reach one node from other, for different $\lambda$ and $\delta$ as in Figure S11.

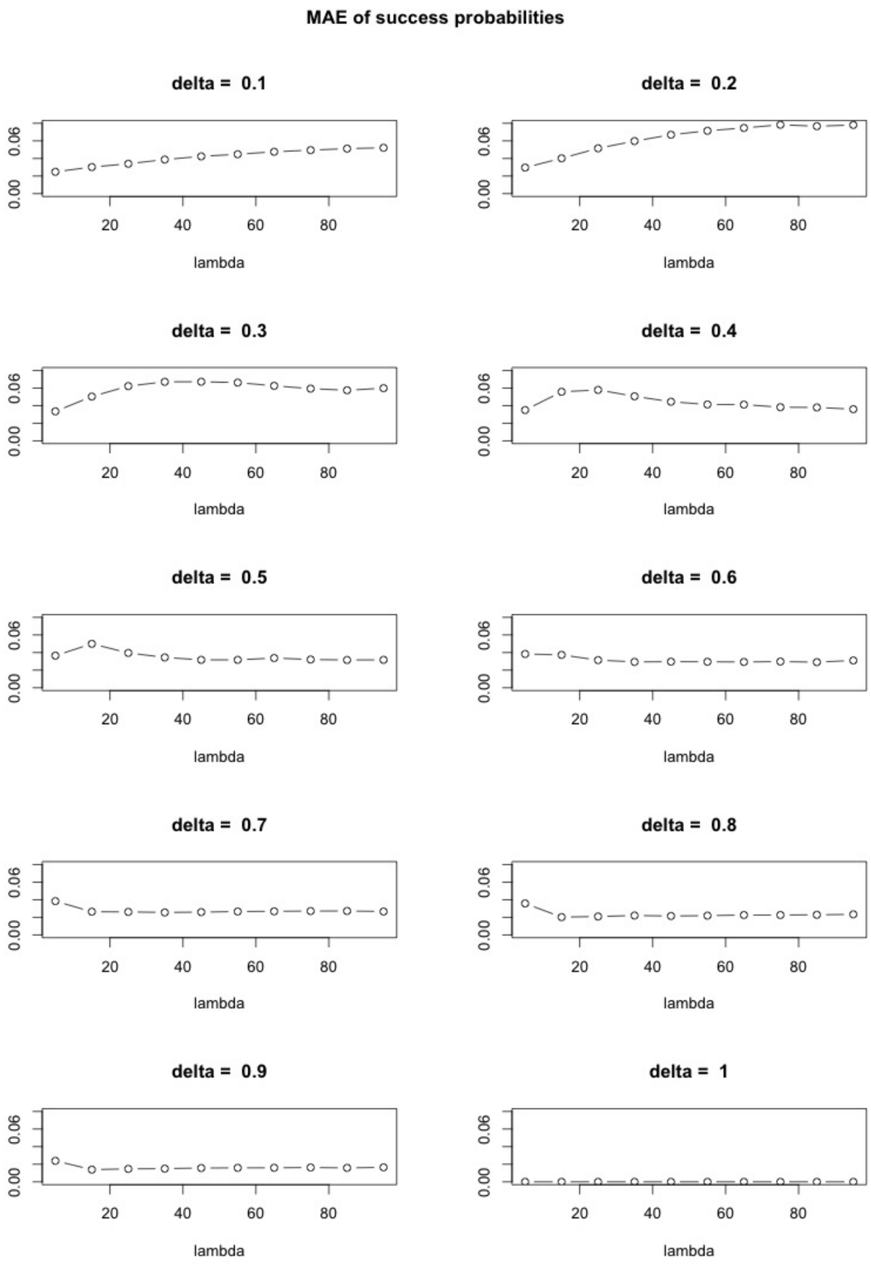


**Fig. S13.** Mean Absolute Error (MAE) of the success probabilities at the tree leaves for different $\lambda$ and $\delta$ as in Figure S11.

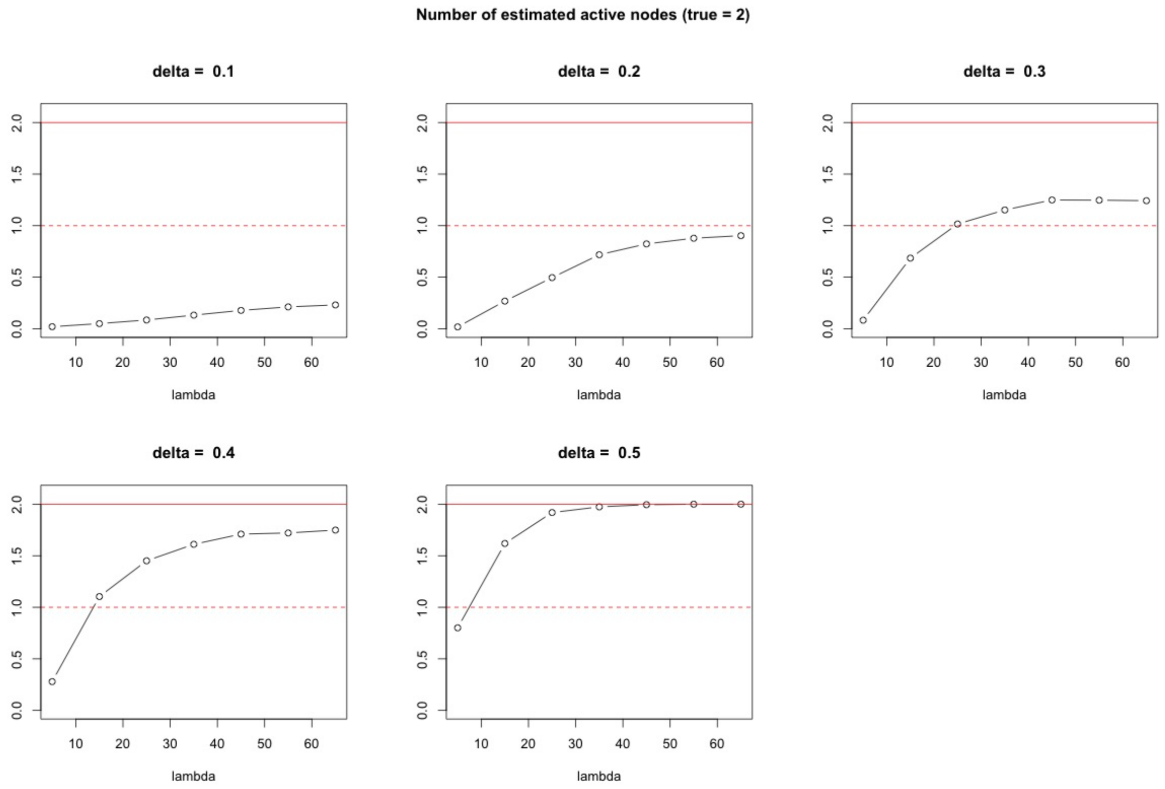


**Fig. S14.** Average number of detected active nodes for different minimal jump sizes $\delta$ and size of active sub-group $\lambda$ in simulations of Setting 3, with true number of active nodes equal to two.

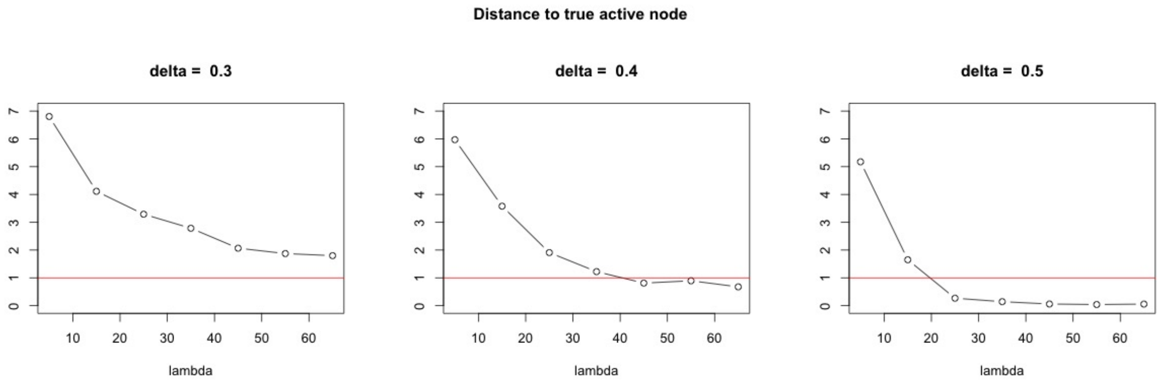


**Fig. S15.** Average distance of some detected active node to first true active node (conditioned on some detection event), with distance function being the minimum number of edges to be traversed to reach one node from other, for different $\lambda$ and $\delta$ as in Figure S14.

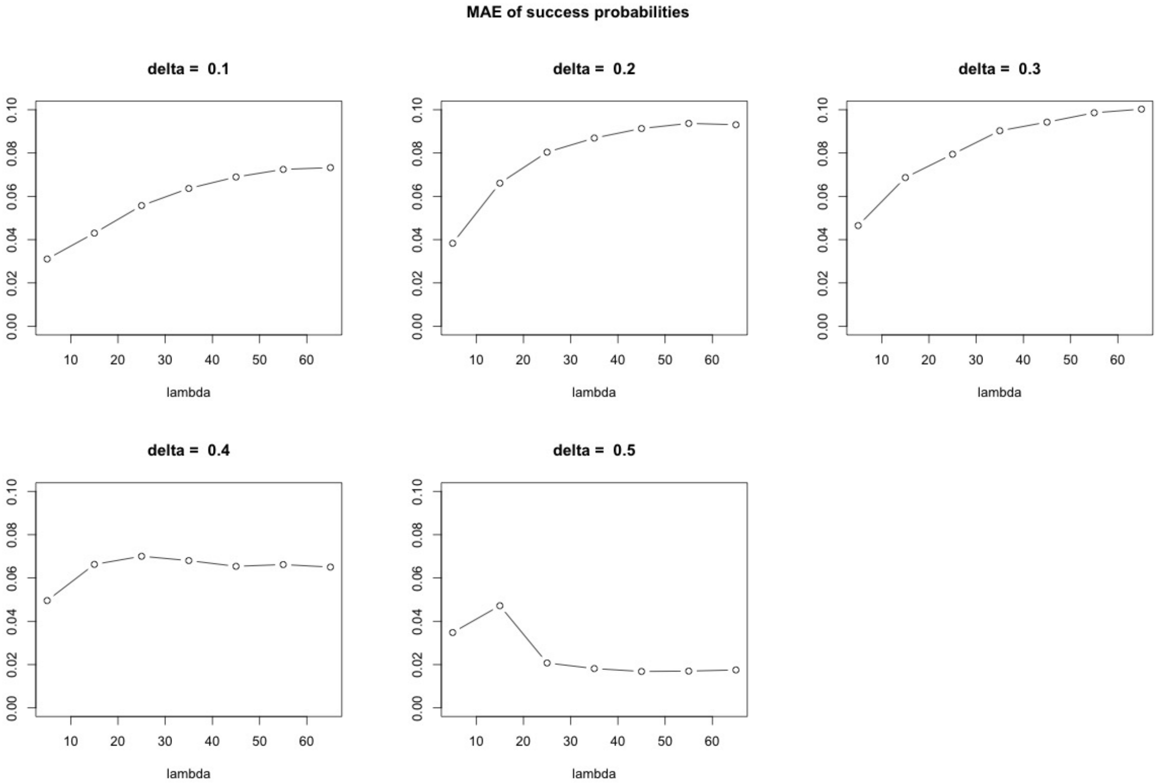


**Fig. S16.** MAE of the success probabilities at the tree leaves for different $\lambda$ and $\delta$ as in Figure S14.

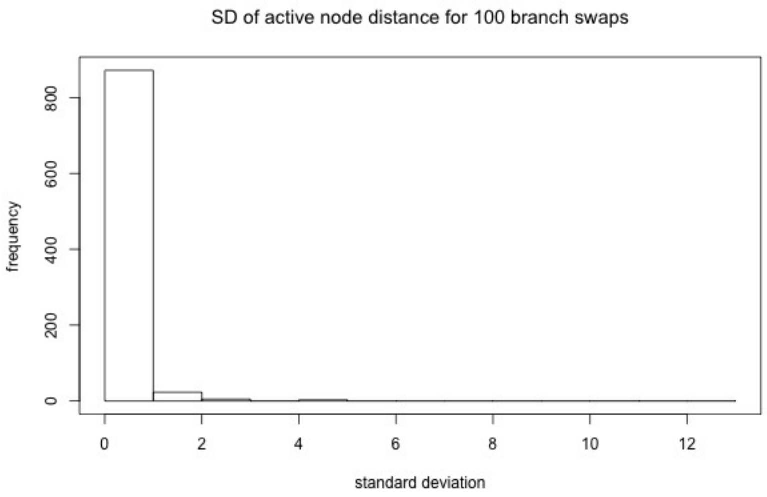


**Fig. S17.** Histogram of $1,000$ samples for the empirical standard deviation of the number of estimated active nodes among $100$ randomly swapped leaf orderings, for reasonable signal to noise ratio with minimal $\lambda = 20$ and $\delta = 0.3$.

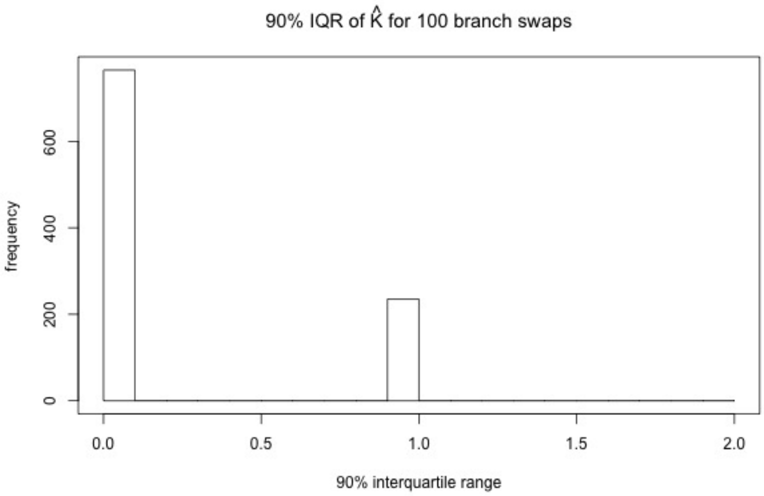


**Fig. S18.** Histogram of $1,000$ samples for the empirical $90\%$ inter quantile distance of the number of estimated active nodes among $100$ randomly swapped leaf orderings, for reasonable signal to noise ratio with minimal $\lambda = 20$ and $\delta = 0.3$.

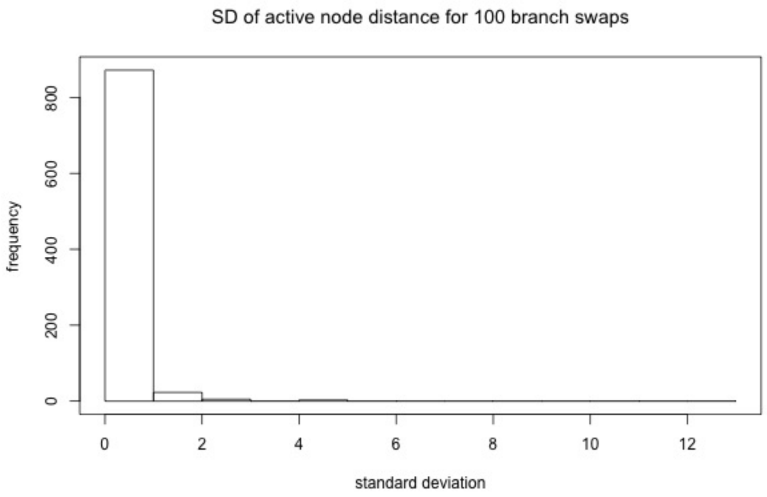


**Fig. S19.** Histogram of $1,000$ samples for the empirical standard deviation of the distance between true and estimated active node among $100$ randomly swapped leaf orderings, for reasonable signal to noise ratio with minimal $\lambda = 20$ and $\delta = 0.3$.

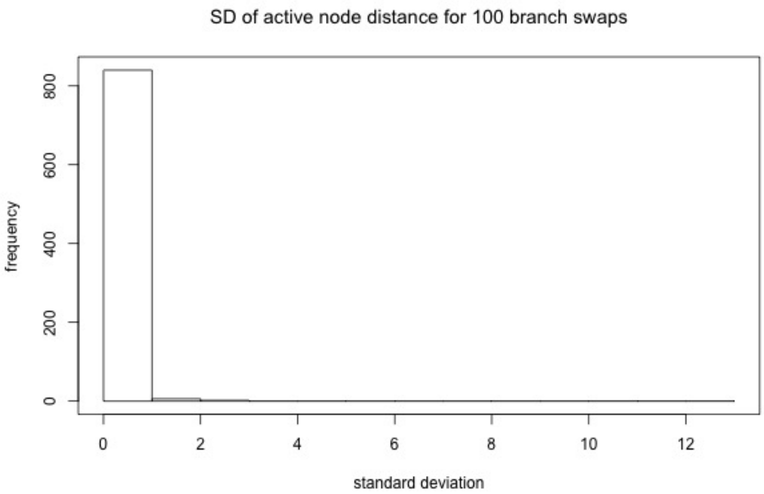


**Fig. S20.** Histogram of $1,000$ samples for the empirical standard deviation of the number of estimated active nodes among $100$ randomly swapped leaf orderings, for high signal to noise ratio regime with minimal $\lambda = 50$ and $\delta = 0.4$.

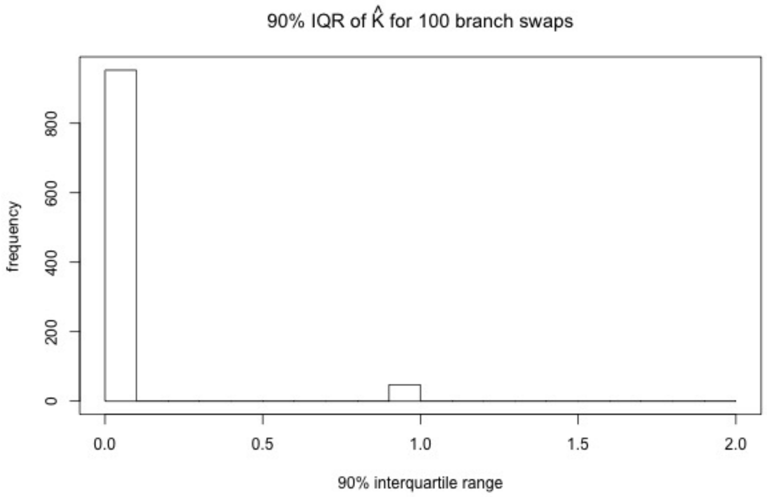


**Fig. S21.** Histogram of $1,000$ samples for the empirical $90\%$ inter quantile distance of the number of estimated active nodes among $100$ randomly swapped leaf orderings, for high signal to noise ratio regime with minimal $\lambda = 50$ and $\delta = 0.4$.

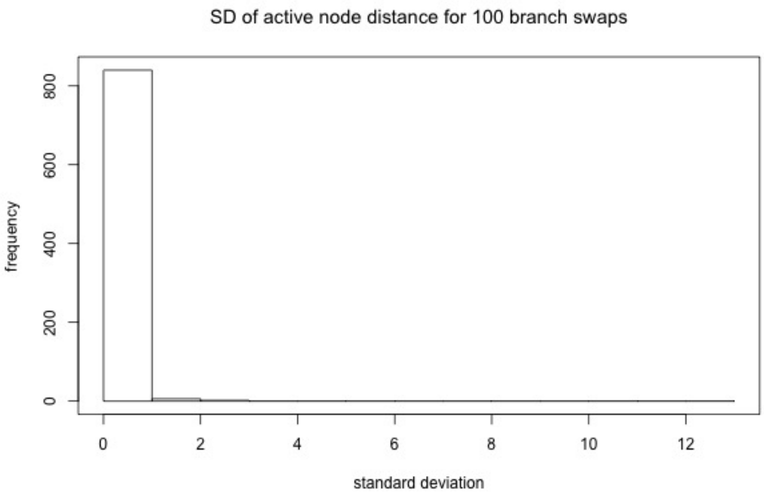


**Fig. S22.** Histogram of $1,000$ samples for the empirical standard deviation of the distance between true and estimated active node among $100$ randomly swapped leaf orderings, for high signal to noise ratio regime with minimal $\lambda = 50$ and $\delta = 0.4$.

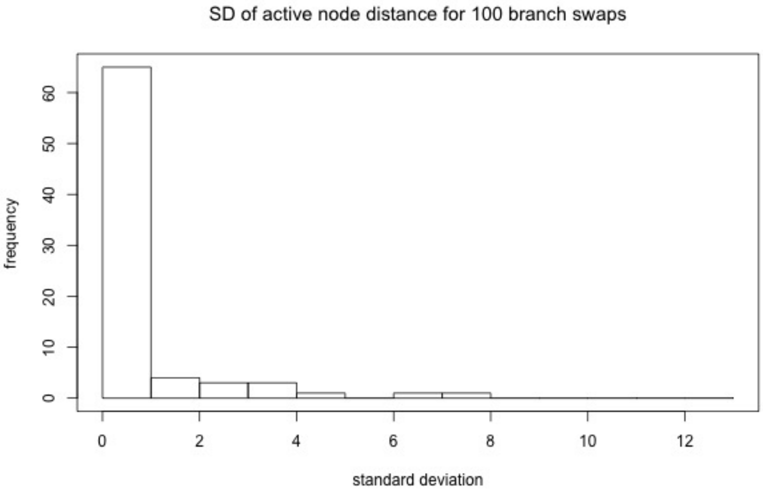


**Fig. S23.** Histogram of $1,000$ samples for the empirical standard deviation of the number of estimated active nodes among $100$ randomly swapped leaf orderings, for low signal to noise ratio with maximal $\lambda = 20$ and $\delta = 0.2$.

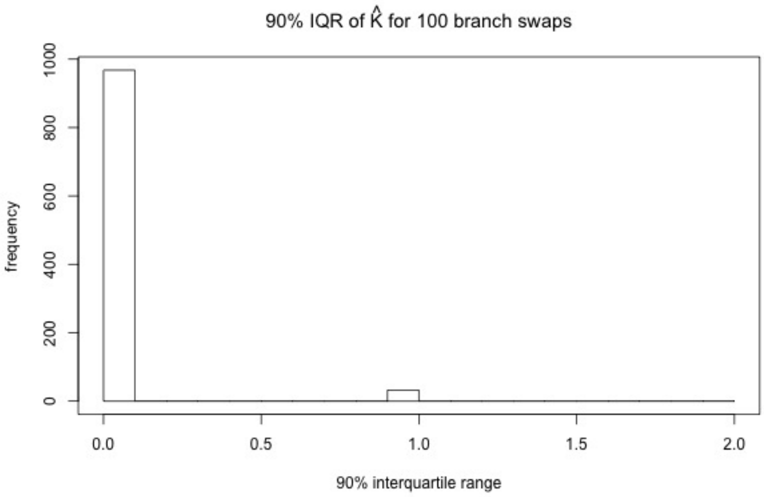


**Fig. S24.** Histogram of $1,000$ samples for the empirical $90\%$ inter quantile distance of the number of estimated active nodes among $100$ randomly swapped leaf orderings, for low signal to noise ratio with maximal $\lambda = 20$ and $\delta = 0.2$.

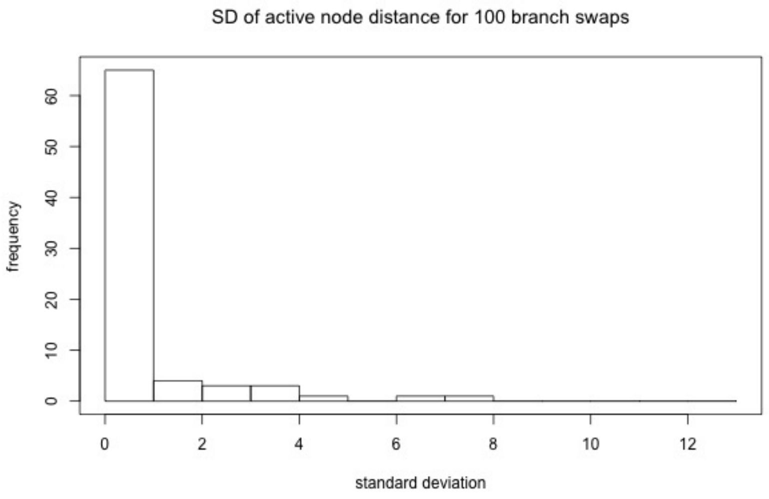


**Fig. S25.** Histogram of $1,000$ samples for the empirical standard deviation of the distance between true and estimated active node among $100$ randomly swapped leaf orderings, for low signal to noise ratio with maximal $\lambda = 20$ and $\delta = 0.2$.

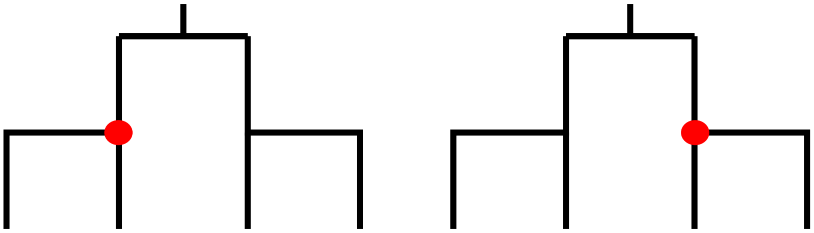

**Fig. S26.** Example of tree with two (left and right) different selections of a single active node (shown in red) which may result in the same vector $(p_i)_{1\leq i\leq n}$ of success probabilities of leaf nodes.

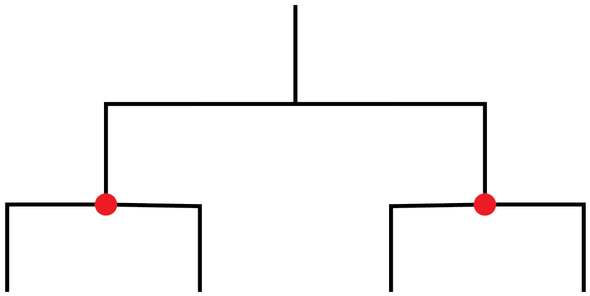

**Fig. S27.** Example of a tree with two active nodes (in red) that are not identifiable, as removing one active node keeps the segmentation unchanged. Note that this is an example with $k = 2$ active nodes but just $2 < k + 1$ different segments and thus, excluded by our analysis.
